# Semantic mining of functional *de novo* genes from a genomic language model

**DOI:** 10.1101/2024.12.17.628962

**Authors:** Aditi T. Merchant, Samuel H. King, Eric Nguyen, Brian L. Hie

## Abstract

Generative genomics models can design increasingly complex biological systems. However, effectively controlling these models to generate novel sequences with desired functions remains a major challenge. Here, we show that Evo, a 7-billion parameter genomic language model, can perform function-guided design that generalizes beyond natural sequences. By learning semantic relationships across multiple genes, Evo enables a genomic “autocomplete” in which a DNA prompt encoding a desired function instructs the model to generate novel DNA sequences that can be mined for similar functions. We term this process “semantic mining,” which, unlike traditional genome mining, can access a sequence landscape unconstrained by discovered evolutionary innovation. We validate this approach by experimentally testing the activity of generated anti-CRISPR proteins and toxin-antitoxin systems, including *de novo* genes with no significant homology to any natural protein. Strikingly, in-context protein design with Evo achieves potent activity and high experimental success rates even in the absence of structural hypotheses, known evolutionary conservation, or task-specific fine-tuning. We then use Evo to autocomplete millions of prompts to produce SynGenome, a first-of-its-kind database containing over 120 billion base pairs of AI-generated genomic sequences that enables semantic mining across many possible functions. The semantic mining paradigm enables functional exploration that ventures beyond the observed evolutionary universe.

## 1. Introduction

While generative AI promises to accelerate the design of functional biological systems, precisely articulating “function” to a generative model is a challenging and often underspecified task. In natural language, the notion of distributional semantics hypothesizes that the meaning of words can be represented by word co-occurrence, i.e., that “you shall know a word by the company it keeps” (Firth, 1957; Harris, 1954) (**Figure 1A**). In biology, an emerging distributional hypothesis defines the function of a gene through its interactions with other genes, i.e., you shall know a gene by the company it keeps (Kwon et al., 2024).

**Figure 1.**
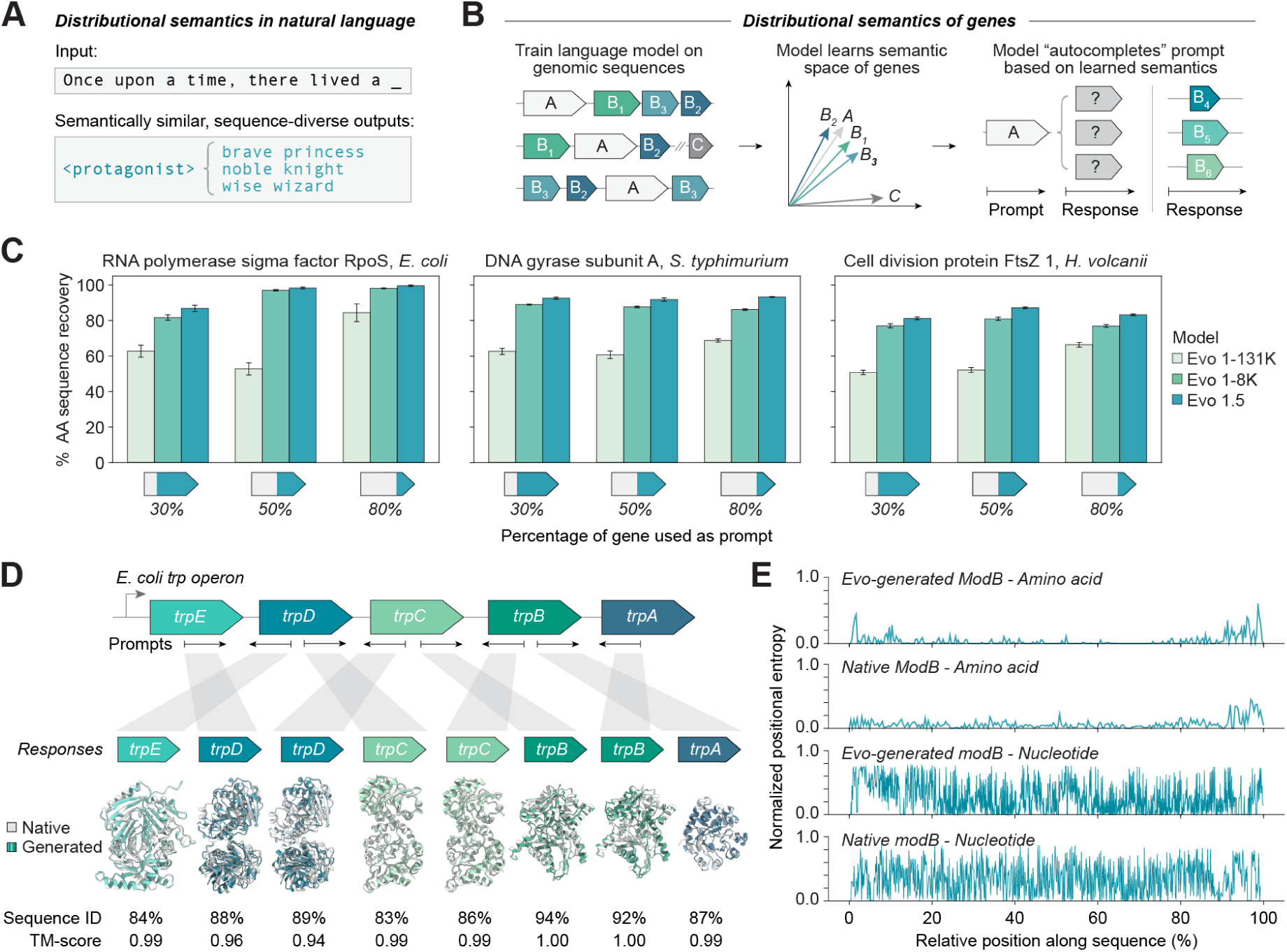
| In-context genomic modeling and design with Evo. **(A)** In natural language, distributional semantics leverages the property that functionally equivalent but lexically distinct words often occupy similar positions in a text with common sets of neighboring words. **(B)** In semantic mining, a genomic language model trained across multiple genes learns to map functionally related genes to similar semantic spaces, enabling generation of functionally related but sequence-diverse genes. **(C)** Sequence recovery assessment when using a genomic language model to autocomplete three conserved prokaryotic genes shows consistent improvement in performance from Evo 1 131K and Evo 1 8K to Evo 1.5, demonstrating enhanced ability to leverage genomic context. AA, amino acid; Bar height, mean; error bars, standard error; *n* = 100. **(D)** Bidirectional completion of conserved *E. coli trp* operon gene sequences demonstrates high sequence identity and structural conservation across the operon. **(E)** Positional entropy comparison between aligned natural and aligned Evo-generated ModB sequences at nucleotide and amino acid levels shows conservation of essential amino acid residues while maintaining high nucleotide diversity. *n* = 500.

In prokaryotic genomes, interacting genes are often positioned directly next to each other on the genome sequence in gene clusters or operons (Jacob and Monod, 1961; Overbeek et al., 1999). Researchers have exploited this property, in a process referred to as genome mining via guilt-by-association, to characterize unknown genes that neighbor functionally characterized genes (Aravind, 2000; Galperin and Koonin, 2000), resulting in the discovery of new molecular mechanisms (Lee et al., 2010; Pers et al., 2015) and important biotechnological tools for genome editing, natural product synthesis, and RNA modification (Medema et al., 2011; Shmakov et al., 2015; Doron et al., 2018; Gao et al., 2020; Blin et al., 2021). At its core, guilt-by-association leverages the distributional hypothesis of gene function to perform function-guided discovery.

A capable generative model of genomic sequences could learn a similar, distributional notion of function to perform function-guided design. Progress in long-context machine learning has enabled advanced generative models of genomic sequences at the multi-kilobase and multi-gene scale (Nguyen et al., 2024). These models are trained to predict the next base pair in a sequence, enabling them to “autocomplete” a partial genomic sequence by generating response sequences to a DNA sequence prompt (**Figure 1B**).

Given the success of guilt-by-association, we reasoned that prompt engineering of a generative genomic model with a sequence of known function could instruct the model to sample novel, functionally related genes in its response. We term this approach “semantic mining,” a new paradigm that harnesses multi-gene relationships to generate novel DNA sequences enriched for targeted biological functions. Notably, unlike traditional genome mining, semantic mining is unconstrained by known evolutionary innovation, potentially allowing us to explore entirely new regions of functional sequence space.

Here, we show that Evo, a 7-billion parameter genomic language model, learns a distributional semantics over genes that enables function-guided design of proteins with high novelty. We first demonstrate that Evo enables in-context genomic design, enabling the successful completion of partial sequences of highly conserved genes and operons. We find that a new version of the previously described Evo model (Nguyen et al., 2024) that we pretrained on 50% more data, which we refer to as Evo 1.5, obtains the best performance on gene-level and operon-level completion tasks.

We then apply semantic mining with Evo to design proteins with both high novelty and specified functional activity. We first focus on the generation of toxin-antitoxin systems, which requires designing two genes that encode a protein-protein interaction. Using a sequential design strategy, we first generate novel toxins by prompting with genomic context around known toxin-antitoxin systems. We then use these Evo-generated toxins as prompts to design their conjugate antitoxin pairs, yielding functional toxin-antitoxin systems containing proteins with remote homology (< 30% sequence identity) to known proteins. We observed that 5 out of 10 designed proteins show antitoxin activity, representing a high experimental success rate of 50%.

To assess Evo’s generalization capability beyond the design of known evolutionary homologs, we then performed semantic mining of systems with high evolutionary novelty. In particular, we prompt-engineered Evo to generate anti-CRISPR proteins that, in nature, have high diversity in both sequence and mechanism and may be produced by *de novo* gene birth (Eitzinger et al., 2020; Huang et al., 2021; Dong et al., 2022). Strikingly, Evo generates anti-CRISPR proteins that effectively inhibit SpCas9-mediated DNA cleavage despite possessing no clear sequence homology to known proteins and low-confidence structure predictions. Evo also designs variants of known anti-CRISPR proteins with stronger inhibitory activity compared to potent natural anti-CRISPRs, as well as an inhibitor with sequence similarity to a protein family not previously associated with CRISPR inhibition. For this task, we also observed a robust experimental success rate of 17%, with 14 out of 84 designs demonstrating anti-CRISPR activity. Together, these results indicate that Evo may be able to generalize beyond the distribution of known natural protein sequences while retaining higher-level, user-defined biological functions.

Given the experimental success of semantic mining in generating novel functional proteins, we then use Evo to generate SynGenome, the first AI-generated genomics database (https://evodesign.org/syngenome/), containing over 120 billion base pairs of synthetic DNA sequences derived from prompts spanning 1,705,667 UniProt entries, 37,108 species, 72,492 protein domains, and 8,914 functional terms. The diversity of coding sequences found in SynGenome recapitulates the natural distribution of protein families, while also containing many generations that go beyond the sequence landscape of the DNA prompts. We make SynGenome and Evo 1.5 openly available to the scientific community to facilitate semantic mining across diverse functions.

In total, Evo can successfully generate proteins with desired functions, often doing so by accessing a highly novel space of sequences that are unlike any known proteins. Semantic mining, with its generalizability and robust success rates, represents a promising new avenue for function-guided generative design. More broadly, we anticipate this work to mark a turning point in biological discovery in which mining of genetic variation becomes substantially augmented by AI-generated sequences that transcend known natural evolutionary constraints.

## 2. Results

### 2.1. Evo enables in-context genomic design

To effectively achieve semantic mining for function-guided design, a model must first understand not just individual gene sequences, but how genes relate to and interact with each other within their broader genomic context. Evo, a 7-billion-parameter genomic language model trained on prokaryotic sequences, is well-positioned to address this challenge through its ability to process long genomic sequences at single-nucleotide resolution (Nguyen et al., 2024).

Similar to how words in language derive meaning from their context (**Figure 1A**), DNA sequences take on functional significance when considered in the context of a gene, an operon, a pathway, or an entire organism. Evo’s efficient long-context architecture can learn how sequence patterns at the level of individual nucleotides relate to a genomic context containing multiple kilobases. Because functionally related sequences cluster together on prokaryotic genomes, supplying an appropriate functional context could condition Evo to generate new genetic sequences with a desired function (**Figure 1B**).

As an initial experiment, we assessed Evo’s ability to leverage genomic context by performing an “autocomplete” task in which we prompted the model with partial sequences of highly conserved prokaryotic genes. We tested three diverse and functionally important genes from bacteria and archaea: RNA polymerase sigma factor *rpoS* from *E. coli*, DNA gyrase subunit A *gyrA* from *S. typhimurium*, and cell division protein *ftsZ1* from *H. volcanii* (Dai and Lutkenhaus, 1991; Chiang and Schellhorn, 2010; Sada et al., 2022) (**Figure 1C**). We also tested three versions of the Evo model: Evo 1 8K, a model pretrained at 8,192 context length; Evo 1 131K, a model made by extending Evo 1 8K to 131,072 context length; and a model newly introduced in this study, which we call Evo 1.5, which extends the pretraining of Evo 1 8K (initially trained on 300 billion tokens) to 450 billion tokens, representing a 50% increase in training data (**Methods**). For each gene, we prompted each Evo model version with varying amounts (30%, 50%, and 80%) of the input sequence and evaluated each model’s ability to complete the remainder of the gene.

Evo 1.5 consistently demonstrated the highest recovery of the native protein sequence, particularly at lower prompt lengths. For instance, with just 30% of the input sequence, Evo 1.5 achieved approximately 85% amino acid sequence recovery for *rpoS*, compared to 65% for Evo 1 131K. This performance advantage was maintained across all tested genes and prompt lengths, with Evo 1.5 achieving near-perfect sequence recovery at 80% input for all targets. These experiments are consistent with previous findings that longer pretraining can improve learning of long-range interactions and high-level concepts in sequence models of natural language and proteins (Lin et al., 2023; Kaplan et al., 2020; Brandes et al., 2022). Moving forward, we therefore selected Evo 1.5 for further investigation and all results attributed to Evo in this study were produced by the Evo 1.5 model.

We further tested Evo’s understanding of genomic context at a multi-gene scale by evaluating its ability to predict gene sequences based on operonic neighbors (**Figures 1D** and **S1**). We prompted the model with sequences of genes either upstream or downstream of target genes in the well-characterized *trp* and *modABC* operons (Maupin-Furlow et al., 1995; Merino et al., 2008), leveraging DNA complementarity to generate sequences bidirectionally. Evo demonstrated robust predictive performance across all tested configurations, achieving over 80% protein sequence recovery for all target genes. Further, the model exhibited adaptability to genomic orientation, successfully generating downstream gene sequences when prompted with reverse complement sequences of upstream genes, and vice versa, while maintaining appropriate directionality across the operon. Notably, even with incomplete sequence recovery, the predicted protein structures of Evo’s generations closely matched the native protein’s conformations, suggesting preservation of functionally critical amino acid residues and structural motifs. These results indicate that Evo not only learns the primary sequence of genes but also captures the broader genomic organization of bacterial operons.

To assess whether Evo’s generations go beyond trivial memorization of training sequences, we analyzed the per-position entropy of both amino acid and nucleotide sequences in the model’s generations (**Figure 1E**). Using the *modABC* operon generations as a test case, we prompted the model with the sequence encoding *modA* from *E. coli* K-12 and analyzed the variability in the generated *modB* responses. The amino acid-level entropy analysis revealed selective conservation, with generally lower entropy at key structural positions and higher variability in less conserved regions. This pattern aligns with natural protein evolution, where functionally critical residues show high conservation while non-essential positions permit amino acid substitutions.

At the nucleotide level, we observed substantially higher entropy across all positions, with variation even in regions coding for conserved amino acids. Given that a single prompt was used to generate all the response sequences, these results suggest that Evo is not simply reproducing memorized sequences but is rather synthesizing information from across its training set to reflect biological constraints while generating diversity in a manner reminiscent of natural evolution.

### 2.2. Semantic mining of functional protein-protein interactions with remote homology to nature

The ability of Evo to understand genomic context consisting of highly conserved genes and operons led us to explore if we could apply semantic mining to less well-conserved biology: phage and bacterial defense systems. These systems, which are shaped by the unrelenting evolutionary arms race between bacteria and phage (Murtazalieva et al., 2024), are some of the most rapidly evolving systems in nature. As a result, defense systems exhibit vast amounts of functional diversity while maintaining minimal sequence conservation across species (Beavogui et al., 2024; Tesson et al., 2024). Interestingly, defense systems of similar functionality also frequently cluster into defense islands, enabling the discovery of novel systems through guilt-by-association approaches (Makarova et al., 2011; Koonin, 2018). To date, some of the most fruitful discoveries of genome mining have been derived from defense systems, including CRISPR-Cas systems, restriction enzymes, and various antimicrobial compounds (Doron et al., 2018; Gao et al., 2020; Picton et al., 2021; Altae-Tran et al., 2023; Cheng et al., 2024). The remarkable success in repurposing these naturally occurring systems for biotechnology applications underscores the vast potential of bacterial defense mechanisms as a source of molecular tools.

Given the natural diversity and genomic co-localization of these systems, we sought to determine if semantic mining could be used to design new defense systems. As an initial test case, we focus specifically on type II toxin-antitoxin (T2TA) systems, some of which play a role in phage defense. Type II toxin-antitoxins consist of a stable toxin protein that can inhibit cell growth or induce cell death under conditions of stress, such as a phage infection, paired with an unstable antitoxin protein that binds and neutralizes the toxin’s activity under homeostatic conditions (**Figures 2A,B**). These systems often maintain conserved high-level genomic architecture despite sequence divergence, with toxin and antitoxin genes typically arranged in adjacent positions (Chan et al., 2023; Guan et al., 2024).

**Figure 2.**
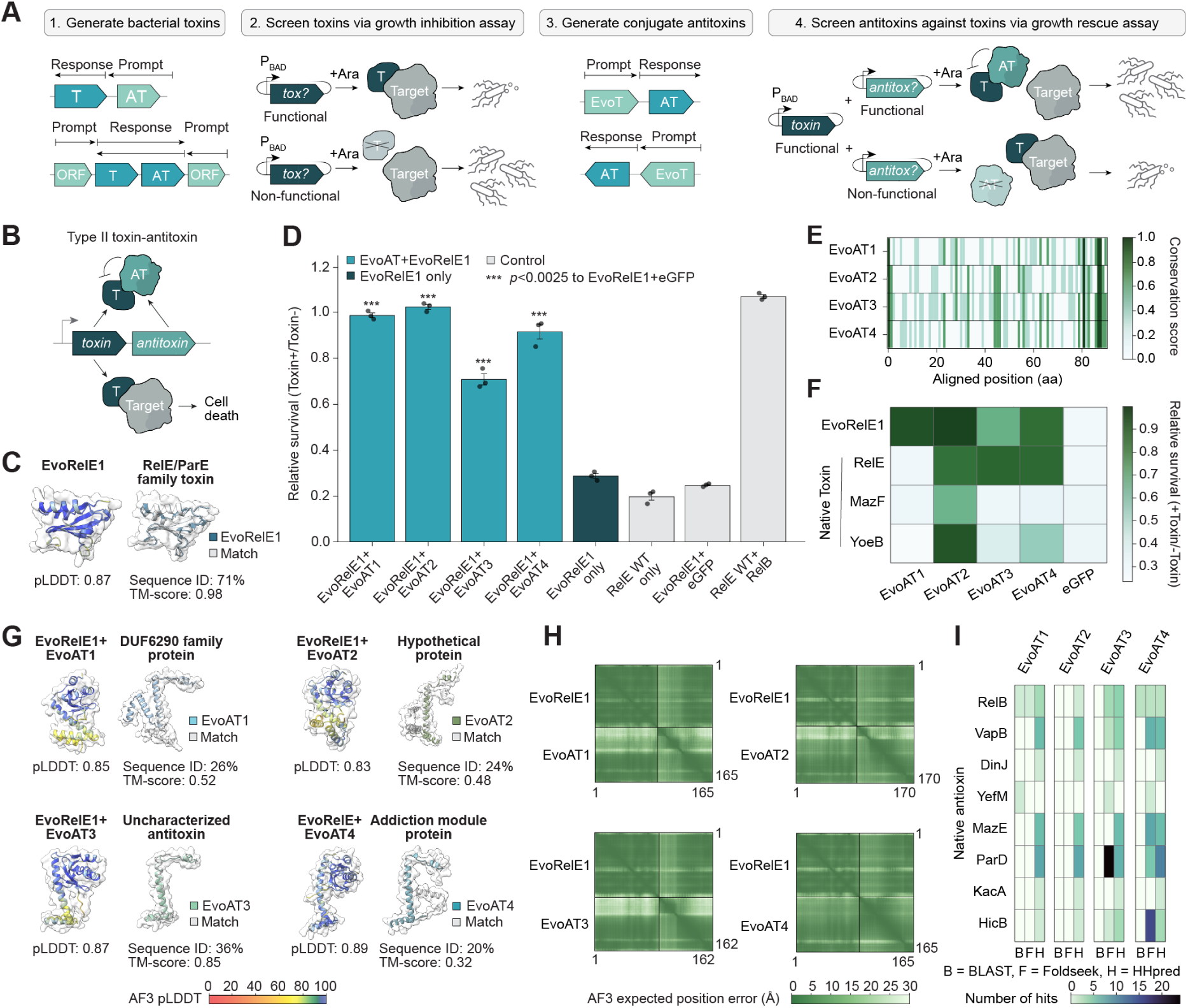
| Evo generates functional toxin-antitoxin protein-protein interactions with remote homology to nature. **(A)** To perform semantic mining of toxin-antitoxin systems, toxins (T) are first generated using toxin genomic context as prompts and tested via growth inhibition assay. Successful toxins (EvoT) then serve as prompts to generate cognate antitoxins (AT), which are finally validated through a growth recovery assay. **(B)** Type II toxin-antitoxin systems function via antitoxin binding and neutralization of toxin activity to prevent cell death. **(C)** AlphaFold 3 structure prediction comparison between functional toxin EvoRelE1 and its closest sequence match. pLDDT, predicted local distance difference test score; TM-score, template modeling score. **(D)** Bar plot showing relative survival rates for different Evo-generated toxin-antitoxin combinations. EvoAT1-4 achieve high growth rescue against EvoRelE1. Bar height, mean; error bars, standard error; circles, biological replicate values; *n* = 3. Statistical significance determined by one-sided Student’s *t*-test comparing to EvoRelE1+eGFP (\*\*\**p* < 0.0025) **(E)** Homology analysis of aligned amino acid sequences for EvoAT1-4, with darker green indicating higher conservation. **(F)** Heat map showing relative survival rates of EvoAT1-4 tested against native toxins. EvoAT2 and EvoAT4 appear to have inhibitory activity against multiple natural toxins. **(G)** AlphaFold 3 structure predictions of EvoAT1-4 in complex with EvoRelE1 and structure comparisons between Evo-generated antitoxins and their closest natural matches. EvoAT1-4 exhibit high confidence predicted structures despite limited sequence homology to their closest protein matches. **(H)** AlphaFold 3 predicted aligned error plots for EvoAT1-4 in complex with EvoRelE1 showing high confidence in predicted structures and interactions. **(I)** Heatmap showing number of structural and sequence matches between Evogenerated antitoxins and known antitoxin families. B, BLAST; F, FoldSeek; H, HHpred.

To generate diversified T2TAs using Evo 1.5, we developed a prompting strategy that leveraged the characteristic co-localization of these systems (**Figure 2A**). In addition to prompting with sequences encoding known T2TAs, we compiled genomic sequences directly upstream and downstream of known T2TA systems. Following sampling using the compiled prompts, we filtered generations for sequences encoding toxin-antitoxin protein pairs that exhibited *in silico* predicted protein complex formation with AlphaFold 3 (Abramson et al., 2024) yet had limited sequence homology to known T2TA proteins.

We first tested Evo-generated toxins using a growth inhibition assay, in which growth arrest indicated successful toxin activity (**Methods**). From these experiments, we were able to identify a functional bacterial toxin, which we call EvoRelE1, that exhibited strong growth inhibition (approximately 70% reduction in relative survival) while possessing 71% sequence identity to a known RelE toxin (**Figures 2C,D**).

We subsequently prompted Evo 1.5 with the sequence of EvoRelE1, hypothesizing that the model could use the genomic context of the toxin sequence to generate a conjugate antitoxin (**Figure 2A**). Using this strategy, we identified a set of 10 antitoxin candidates exhibiting little to no sequence homology to natural protein sequences that were tested for their ability to block the toxin activity of EvoRelE1.

Following co-expression with EvoRelE1, half of the Evo-generated antitoxin candidates were able to rescue cell growth (**Figures 2D** and **S2**). The most effective candidates, designated as EvoAT1 and EvoAT2, restored growth to 95-100% of normal cell survival, with candidates EvoAT3 and EvoAT4 demonstrating moderate rescue activity (70% and 90% relative survival, respectively). Sequence analysis of the successful antitoxins revealed discrete regions of conservation (**Figure 2E**), potentially highlighting key motifs required for toxin neutralization despite their overall sequence diversity. Furthermore, when tested against native RelE, MazF, and YoeB toxin systems, several of the Evo-generated antitoxins were able to rescue growth, with EvoAT2 showing inhibitory activity against all three toxins and EvoAT4 rescuing growth against native RelE and YoeB toxins (**Figure 2F**). This observation suggests that the EvoATs may be able to inhibit multiple toxins despite sharing limited sequence homology with their natural antitoxin counterparts (**Figures 2F** and **S3**). Interestingly, when co-folded using *in silico* structure prediction methods, several of the EvoATs had low-confidence predicted complex formation with the natural toxins that they inhibited (EvoAT4 + YoeB ipTM = 0.14, EvoAT2 + YoeB ipTM = 0.47, EvoAT2 + MazF ipTM = 0.34) (**Figure S4**), highlighting the potential for semantic mining to identify novel molecular interactions that extend beyond the domain of state-of-the-art structure prediction models.

EvoAT1 through EvoAT4 all have sequence identities with natural proteins that fall near the “twilight zone” of sequence similarity (20% to 36%), a range where sequence similarity alone is insufficient to reliably predict shared structure or function (Rost, 1999). Both EvoAT1 and EvoAT2 only exhibit homology to proteins not actively annotated as antitoxins, showing just 26% sequence identity to a DUF6290 family protein from *Actinomyces* and 24% sequence identity to an uncharacterized *Magnetococcus sp*. YQC-5 hypothetical protein respectively (**Figure 2G**). Given that EvoAT2’s closest match is an uncharacterized hypothetical protein that appears in a similar genomic context (**Figure S5**), our results suggest this protein may function as part of a toxin-antitoxin system. Similarly, the homology between EvoAT1 and the DUF6290 family reinforces a hypothesized potential role for this domain of unknown function in antitoxin activity (Blum et al., 2024). These findings suggest that Evo’s understanding of genomic context enables functional annotation of previously uncharacterized proteins, as well as function-conditioned design that is unconstrained by existing annotations. The structural predictions for EvoAT1 and EvoAT2 revealed high-confidence folds (pLDDT scores of 0.85 and 0.83, respectively) (**Figure 2G**), and when co-folded with EvoRelE1, both antitoxins exhibited minimal predicted position error (**Figure 2H**). These structural characteristics and high functional activity are particularly noteworthy given the antitoxins’ limited sequence homology to known antitoxin proteins (**Figure 2I**). Overall, these results further underscore the ability of Evo to generate functional proteins with remote homology to natural proteins.

Sequence and structural homology searches using BLAST, HHpred, and Foldseek (van Kempen et al., 2024; Söding et al., 2005; Gabler et al., 2020; Zimmermann et al., 2018; Sayers et al., 2022) revealed that the EvoATs show similarity to multiple independent antitoxin superfamilies, particularly ParD, MazE, and VapB (**Figure 2I**). This finding, coupled with the activity of EvoAT2 and EvoAT4 against multiple toxins, is especially notable because their cognate toxins employ fundamentally different mechanisms of action. In natural systems, these mechanistic differences have typically led to the evolution of distinct antitoxin structures specialized for neutralizing each type of toxin (Fraikin et al., 2020; Qiu et al., 2022). This suggests that Evo may have identified a broader functional compatibility between antitoxin and toxin families than previously recognized, highlighting the potential of semantic mining to reveal new insights into protein-protein interaction patterns in prokaryotic systems.

### 2.3. Semantic mining of functional *de novo* genes

Given the experimental success of Evo-generated functional proteins with remote homology to nature, we next explored whether semantic mining could learn to generate even greater evolutionary novelty. Anti-CRISPRs (Acrs) are proteins used by phages to neutralize bacterial CRISPR-Cas immune systems. Given their critical role in helping phages evade CRISPR-Cas mediated cleavage (**Figure 3A**), Acrs represent one of the most striking examples of rapid protein evolution in bacterial-phage arms races, with many Acrs appearing to be novel innovations lacking detectable homology to other protein families, including other Acrs (Davidson et al., 2020; Eitzinger et al., 2020; Huang et al., 2021). This remarkable diversity is reflected in their varied mechanisms of action, from direct CRISPR-Cas protein binding to DNA mimicry to transcriptional silencing (Shin et al., 2017; Camara-Wilpert et al., 2023; Choudhary et al., 2023; Trost et al., 2024), making them valuable tools for understanding protein evolution and developing synthetic control systems for CRISPR technologies (Pawluk et al., 2018).

**Figure 3.**
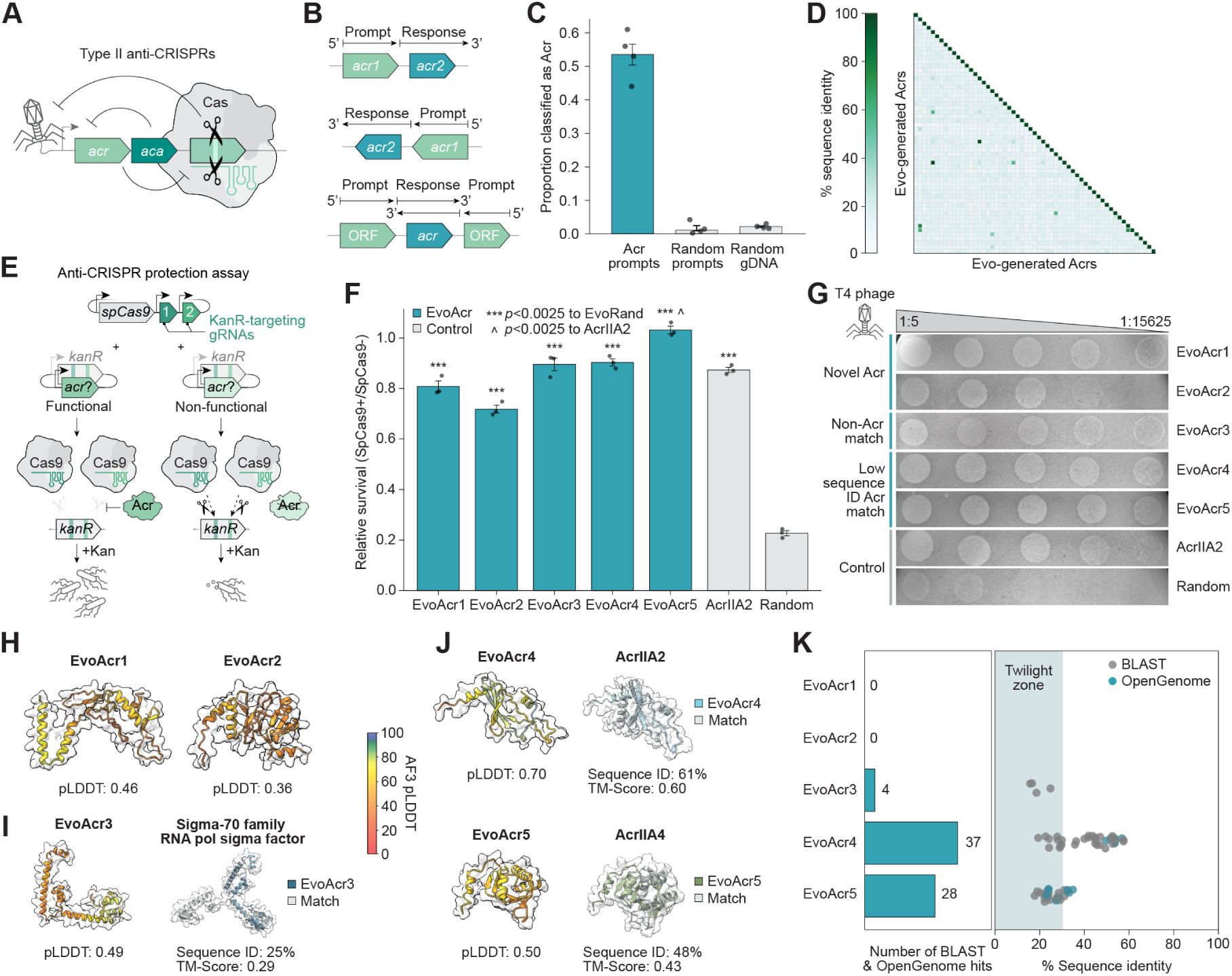
| Evo generates functional anti-CRISPR proteins with no homology to known proteins. **(A)** Type II anti-CRISPR systems involve an anti-CRISPR (*acr*) gene that encodes an Acr protein that inhibits type II-A Cas nuclease activity, often co-occurring with anti-CRISPR associated genes (*aca*). **(B)** To perform semantic mining of Acrs, known genomic contexts of type II anti-CRISPRs, including potential *aca* genes, are used as prompts. **(C)** PaCRISPR classification demonstrates significant enrichment of Acr-like sequences in generations from Acr prompts compared to controls. Bar height, mean; error bars, standard error; circles, different batches of generations; *n* = 4. **(D)** Sequence identity matrix demonstrating high diversity among random sample of Evo-generated Acrs. **(E)** In the anti-CRISPR protection assay, functional Acr proteins prevent SpCas9-mediated cleavage of a kanamycin resistance gene, enabling cell survival in kanamycin-supplemented media, while nonfunctional candidates allow SpCas9 targeting and cell death. **(F)** Bar plot showing relative survival rates for Evo-generated Acrs. EvoAcr1-5 achieve robust protection against SpCas9, with EvoAcr5 exceeding the activity of the AcrIIA2 positive control. Bar height, mean; error bars, standard error; circles, biological replicate values; *n* = 3. Statistical significance is determined by a one-sided Student’s t-test compared to random control *(*** p < 0.0025)* and AcrIIA2 (^*p* < 0.0025). **(G)** T4 phage plaque assays validating anti-CRISPR activity. *E. coli* cells co-expressing SpCas9 targeted to T4 phage and candidate Acrs were embedded in soft agar, and serial dilutions of T4 phage were spotted onto the bacterial lawn. Plaque formation indicates successful anti-CRISPR protection. Experiment performed in triplicate with representative images shown. **(H)** AlphaFold 3 structure predictions for EvoAcr1-2 showing low confidence predictions. pLDDT, predicted local distance difference test score; TM-score, template modeling score. **(I)** AlphaFold 3 predicted structure of EvoAcr3 and structure alignment between EvoAcr3 and its closest sequence match, revealing limited homology to a sigma-70 family protein. **(J)** AlphaFold 3 structure predictions comparing EvoAcr4-5 with their closest Acr sequence matches, showing moderate structural and sequence similarity. **(K)** Sequence similarity analysis of EvoAcr1-5 calculated by BLAST search against BLAST-nr and MMseqs2 search against OpenGenome. EvoAcr1-2 demonstrate no significant homology to known proteins, EvoAcr3 shows matches only in the “twilight zone” of sequence similarity (Rost, 1999), and EvoAcr4-5 show limited sequence matches to known Acrs.

Despite their sequence diversity, many Acr operons maintain a somewhat conserved operonic architecture, consisting of multiple Acr genes appearing together alongside neighboring anti-CRISPR associated (*aca*) genes (Stanley et al., 2019) (**Figure 3B**). This architectural conservation, combined with their frequent emergence as *de novo* genes, makes Acrs an ideal test case for assessing semantic mining’s ability to generalize with respect to sequence and structure while retaining a desired higher-level function.

Using genomic sequences from known Cas9-targeting Acr genes and their associated operons as prompts, we used Evo 1.5 to generate 3,160 sequences totaling 3.16 million base pairs. After filtering for size, complexity, and secondary structure, we identified 468 potential Acr open reading frames (ORFs). To handle the high sequence diversity of Acr proteins complicating traditional MSA and structural filtering methods, we next used PaCRISPR, a machine learning model trained to identify potential Acr proteins, to evaluate our generated candidates (Wang et al., 2020). Consistent with our prompts providing successful functional conditioning, generations derived from Acr-containing genomic contexts were substantially more likely to be classified as potential Acrs by PaCRISPR compared to negative control sequences (**Figure 3C**). Approximately 55% of minimally filtered proteins generated from Acr-associated prompts were classified as likely Acrs, compared to less than 5% of proteins generated from either random prompts or random genomic sequences. Further, the distribution of sequence identities among the predicted Acrs showed a wide range of novelty, with most candidates showing low similarity to each other (**Figure 3D**) (median pairwise sequence identity = 23%). This enrichment for diverse Acr-like sequences demonstrates that semantic mining can effectively bias generations toward a desired function by leveraging genomic context alone, even in the absence of clear sequence or structural signatures.

To test the Acr candidates’ protection ability against SpCas9, we co-transformed *E. coli* with plasmids encoding our candidate Acrs and a CRISPR-targeted kanamycin resistance gene, where functional Acr protection would enable bacterial growth in kanamycin-supplemented media through inhibition of CRISPR-mediated cleavage (Forsberg et al., 2019) (**Figure 3E**). From this initial experiment, we found that 17% of tested sequences exhibited some degree of anti-CRISPR activity (**Figure S6**), a high success rate when considering the lack of structure or sequence homology-guided design. From this pool, we further identified five promising proteins (EvoAcr1-5) that demonstrated strong protection against SpCas9-mediated targeting in both liquid culture survival assays (**Figure 3F**) and phage infection experiments (**Figures 3G** and **S7**).

Detailed bioinformatic analysis of these five Acrs revealed a high level of diversity in their sequence origins. EvoAcr4 and EvoAcr5 share moderate sequence similarity to known Acrs (61% and 48% to AcrIIA2 and AcrIIA4, respectively) and demonstrated robust protection against SpCas9, with EvoAcr5 showing activity exceeding that of the positive control AcrIIA2 in liquid culture assays (relative survival rates of 1.01 to 0.87 respectively) (**Figures 3F,H**). EvoAcr3 presented an intriguing case: while sharing only “twilight zone” sequence identity (Rost, 1999) and limited structural alignment (sequence ID = 25%, TM-score = 0.29) with a sigma-70 family RNA polymerase sigma factor, it maintained strong anti-CRISPR activity (relative survival of 0.89) (**Figures 3F,I**). This suggests a potential mode of CRISPR inhibition that is not described by existing functional databases.

Most notably, EvoAcr1 and EvoAcr2 represented proteins that eluded both sequence and structural homology characterization, showing no significant identity to proteins in OpenGenome (the Evo training data) or to known proteins in the nr database by BLAST (**Figures 3J,K**). The AlphaFold 3-predicted structures of EvoAcr1 and EvoAcr2 also have low confidence pLDDT scores of 0.46 and 0.36, respectively (**Figures 3H** and **S8**). Despite this lack of sequence or structural similarity to known proteins, they demonstrated robust protection in both liquid culture and phage infection assays, with relative survival rates of 0.82 and 0.74, respectively. This experimental validation of novel, functional Acrs supports the ability of semantic mining to access entirely unexplored regions of sequence space while maintaining specific desired functions. Together with EvoAcr3-5, these results demonstrate that genomic context can guide the generation of diverse anti-CRISPR proteins, from sequence variants of known architectures to entirely new protein sequences.

### 2.4. SynGenome: A first-of-its-kind synthetic genomic database

Following our experimental validation that semantic mining can generate novel, functional proteins from genomic context alone, we reasoned that this approach could be systematically applied to generate functional genes across prokaryotic biology. To this end, we developed SynGenome, a comprehensive database of synthetic DNA sequences generated using Evo. In line with the function-guided design principles underlying semantic mining, we prompted the model with sequences derived from over 1.5 million prokaryotic and phage proteins to generate a large set of diverse sequences that could be enriched for novel genes functionally related to the prompt.

To construct SynGenome, we leveraged the UniProt database to identify protein-coding genes and their adjacent sequences from prokaryotic organisms and bacteriophages. For each coding sequence, we generated six distinct prompts: the upstream region, coding sequence (CDS), downstream region, and their respective reverse complements (**Figure 4A**). Using the Evo 1.5 model with standard autoregressive decoding, we generated multiple synthetic sequences for each prompt, resulting in a database containing over 120 billion DNA base pairs (**Methods**).

**Figure 4.**
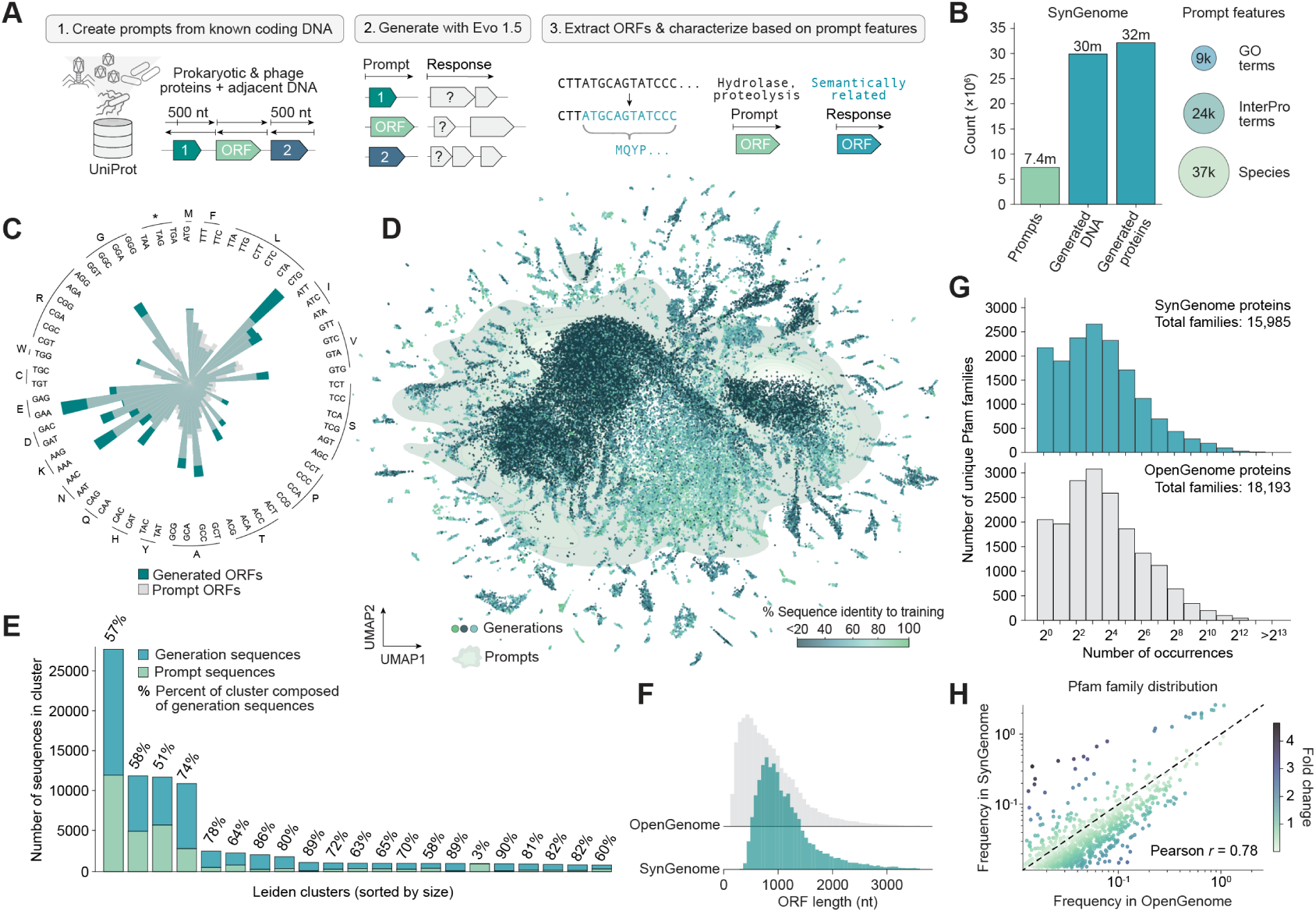
| 120 billion base pairs of AI-generated genomic sequences with SynGenome. **(A)** To construct SynGenome, we created prompts from known-protein coding genes, generated synthetic DNA sequences using Evo, and bioinformatically characterized the generated sequences. **(B)** Number of prompts and generated sequences in SynGenome, and their associated features. GO, gene ontology. **(C)** Codon usage bias of Prodigal-predicted ORFs in prompt sequences and generated sequences, demonstrating preservation of codon preferences. **(D)** UMAP of jointly projected Evo embeddings of SynGenome and prompt sequences. Generation embeddings are colored by sequence identity to training data and prompt embeddings are shown in light green. **(E)** Bar plot showing relative proportions of generation and prompt sequences in most populous Leiden clusters, with percentages indicating fraction of generated sequences per cluster. **(F)** Prodigal-predicted ORF lengths in for 5000 nt representative samples of OpenGenome and SynGenome, with ORFs showing similar length distributions. nt, nucleotide. **(G)** Histogram showing number of Pfam protein family occurrences in SynGenome and OpenGenome, with both following a similar long-tailed distribution of family abundance. **(H)** Scatterplot showing correlation between protein family frequencies in SynGenome versus OpenGenome, demonstrating preservation of natural protein family abundance patterns.

To facilitate functional exploration of the database, we organized the synthetic sequences according to the Gene Ontology (GO) terms and InterPro domain annotations associated with their corresponding prompts (**Figure 4B**), as the generated sequences are likely to be enriched for functionally related elements. As generating sequences with language models can often be prone to repetitive generations, we implemented minimal filtering to remove highly repetitive low complexity sequences, for example, long stretches of sequence containing only a single base pair (**Methods**).

To characterize the generations in SynGenome, we first examined codon usage patterns between the generated sequences and the prompts. This analysis revealed that generated sequences closely mirrored their prompts, with the synthetic sequences maintaining similar codon preferences to natural coding sequences (**Figure 4C**). We further examined the relationship between prompts and their corresponding generated sequences in Evo embedding space (**Figure 4D**) by performing Leiden clustering and analyzing the relative proportions of each sequence type within clusters (**Figures 4E** and **S9**). Most Leiden clusters contain a mixture of generated sequences and natural prompt sequences, indicating that many generations occupy similar regions of the language model embedding space. However, we observed that 54 clusters representing 19% of the generated sequences are almost completely composed of generated sequences, potentially indicating regions of synthetic sequences that extend beyond the semantic space represented by natural sequence embeddings.

When compared to sequences from OpenGenome, we found that SynGenome-generated ORFs closely mirrored the natural prokaryotic ORF length distribution predicted by Prodigal (Hyatt et al., 2010) (**Figure 4F**). At the protein level, SynGenome matched both the global distribution of Pfam domain frequencies found in natural proteins (**Figure 4G**), with specific Pfam families also showing similar occurrence frequencies between synthetic and natural sequences (Mistry et al., 2021; Blum et al., 2024) (Pearson correlation coefficient, *r* = 0.78, (**Figure 4H**). These analyses demonstrate that SynGenome recapitulates the architecture and functional diversity of natural prokaryotic sequences.

Collectively, these results highlight the potential for SynGenome to become a valuable tool for exploring and expanding protein function through semantic relationships. By mining SynGenome, researchers could discover functionally related proteins that extend beyond natural sequence space, gain insights into potential functions of uncharacterized proteins based on genetic co-occurrence patterns, and create diverse screening libraries for engineering desired functions (**Figure S10**).

SynGenome is freely available as a public resource at https://evodesign.org/syngenome/, where researchers can search the database using protein names, UniProt IDs, InterPro domains, species names, or GO terms of interest. This allows for rapid identification of relevant synthetic sequences, enabling researchers to quickly find and explore generations that align with their research interests. We anticipate that SynGenome can serve as a practical tool that facilitates gene discovery and engineering with semantic mining for the broader scientific community.

## 3. Discussion

Advanced genomic sequence models, trained on hundreds of billions of DNA base pairs across the diversity of prokaryotic life, can enable unprecedented capabilities. We demonstrate that Evo enables controllable design of desired functions when prompted with sequences that are meant to elicit those functions, with high experimental success rates of 17-50% when testing only tens of variants in the laboratory. Many of the designed proteins also have no significant homology to proteins of similar function or, in some cases, to any known protein. These results blur the line between “*de novo*” protein design (Huang et al., 2016; Watson et al., 2023; Kortemme, 2024) and diversification based on evolutionary models (Madani et al., 2023; Hayes et al., 2024; Hie et al., 2024), providing an “existence proof” that sequence models without any structural conditioning can meaningfully generalize beyond natural sequence space.

Semantic mining represents a fundamentally new paradigm for protein design that is highly complementary to existing approaches. First, unlike design methods based on task-specific fine-tuning (Madani et al., 2023; Jiang et al., 2024; Nguyen et al., 2024), semantic mining requires no additional training or supervision that could bias generations toward the distribution of known examples. Second, in contrast with approaches that propose specifying function through natural language descriptions found in existing knowledgebases (Praljak et al., 2024), semantic mining accesses the vast functional diversity embedded within genomic sequences. This enables functional design that can leverage biological processes that are not yet documented in the scientific literature. For example, we generate antitoxins that suggest broader functional compatibility between diverse toxin and antitoxins systems than previously recognized (**Figure 2F**) and a protein with anti-CRISPR activity that maps to a protein family with a different putative function (**Figure 3I**). Third, by leveraging genomic context as functional conditioning, semantic mining does not require existing structural or mechanistic hypotheses; indeed, protein design pipelines that filter out low-confidence structure predictions (Verkuil et al., 2022; Pacesa et al., 2024) would have removed most of our high-activity designs. Genomic conditioning is also useful when specifying functions such as anti-CRISPR activity that could be accomplished by many structures and mechanisms (Choudhary et al., 2023). Semantic mining of genomic language models therefore represents a powerful and orthogonal approach to current biological design strategies.

Traditional genome mining via guilt-by-association, which motivates many of the ideas in this study, is constrained to discovered evolutionary diversity that was generated over billions of years. In contrast, semantic mining enables the generation of a near limitless supply of sequence diversity for a biological system of interest. To facilitate broader accessibility of this new source of evolutionary material, we report SynGenome, a database of over 120 billion base pairs of AI-generated genomic sequences, which we make publicly available. This resource enables researchers, especially those without the computing resources required to conduct large scale sampling from a generative model, to mine for sequences related to their function of interest. This data could potentially contain new molecular tools and provide insights into protein function and evolution.

While semantic mining represents a new level of sequence novelty and functional improvement for generative genomics, several fundamental limitations and challenges remain. Autoregressive generation is prone to sampling repetitive sequences or to hallucinating realistic but non-functional designs. Semantic mining therefore requires both *in silico* filtering and experimental testing to validate downstream functions. Semantic mining is also theoretically limited to functions that are found in nature, particularly those that are encoded in prokaryotic organisms. However, we note that only a small fraction of this functional diversity has been characterized by the biological literature and that mining of prokaryotic functional diversity has directly led to powerful biotechnologies such as polymerase chain reaction (PCR), optogenetics, genome editing, and genetic recombination (Chien et al., 1976; Nagel et al., 2003; Boyden et al., 2005; Govorunova et al., 2017; Altae-Tran et al., 2023; Durrant et al., 2024).

Looking forward, increased scaling of pretrained models and availability of training data from rapidly growing sequencing resources will likely reinforce the capabilities of semantic mining. We also anticipate combining the rich information learned by pretrained models with more advanced inference-time strategies will improve generation quality. Analogous to generative models of images that exhibit improved controllability when given prompts that are themselves generated by a natural language model (Esfandiarpoor and Bach, 2024; Mañas et al., 2024), synthetic prompts could be used to improve the generation quality of genomic language models as well; for example, EvoAT1-4 were all generated in the context of the previously designed EvoRelE1. Further, given the utility of chain-of-thought prompting in natural language (Wei et al., 2023), genomic language models that sequentially generate multicomponent systems could accelerate the development of synthetic biological circuits, molecular machines, metabolic pathways, or even complete genomes. Lastly, techniques for mechanistic interpretability of large language models (Bricken et al., 2023; Templeton et al, 2024) or post-training that leverages high-throughput experimental data collection (Ouyang et al., 2022; Rafailov et al., 2024; Rai et al., 2024) could improve the quality and diversity of generated functions. By leveraging semantic mining, exploration of synthetic genomic space may reveal biological discoveries that complement and extend beyond those discovered in natural organisms.

## 4. Code availability

Code for generation with Evo based on a user-defined prompt is available at https://github.com/evo-design/evo/. Scripts for semantic mining of toxin-antitoxin systems and anti-CRISPRs are available https://github.com/evo-design/evo/tree/main/semantic_mining. The Evo 1.5 model is available under an open source license at Hugging Face: https://huggingface.co/evo-design/evo-1.5-8k-base.

## 5. Data availability

SynGenome is explorable and searchable at https://evodesign.org/syngenome/. Raw data has been deposited to Hugging Face datasets at https://huggingface.co/datasets/evo-design/syngenome-uniprot.

## 6. Materials availability

DNA, RNA, and protein sequences used during our validation experiments are available in **Data S1**. All newly created materials are available upon reasonable request to the corresponding authors.

## 7. Acknowledgements

We thank Kevin Forsberg, Regan Russell, and Samantha Sakells for supplying plasmid backbones and for helpful discussions on anti-CRISPR experiments. We thank Matthew Durrant, Patrick Hsu, Julia Kazaks, David Li, Talal Widatalla, Brandon Ameglio, Sergey Ovchinnikov, April Pawluk, Brian Plosky, and Chiara Ricci-Tam for helpful discussions and assistance with manuscript preparation. A.T.M. acknowledges funding support from the Knight-Hennessy Graduate Scholarship Fund. A.T.M. and S.H.K. acknowledge funding support from the National Science Foundation Graduate Research Fellowship Program. B.L.H. acknowledges funding support from Arc Institute, Stanford Institute for Human-Centered Artificial Intelligence (HAI) Hoffman-Yee Research Grants, V. Gupta, and R. Tonsing.

## 8. Author Contributions

A.T.M., S.H.K., and B.L.H. conceived the project. B.L.H. supervised the project. E.N. and B.L.H. trained Evo 1.5. A.T.M. developed code for and performed the gene completion, operon completion, and entropy analyses. A.T.M. and S.H.K. compiled prompts for the Acr sampling. A.T.M developed code for and performed the analysis and filtering of the Acr candidates. A.T.M and S.H.K. experimentally tested the Acr generations. A.T.M compiled prompts for the toxin-antitoxin sampling. A.T.M developed code for and performed the analysis and filtering of the toxin-antitoxin candidates. A.T.M experimentally tested the toxin-antitoxin generations. A.T.M compiled the prompts for SynGenome. A.T.M and B.L.H. developed the code for and performed the sampling for SynGenome. B.L.H. developed the SynGenome website. A.T.M. S.H.K., and B.L.H. wrote the first draft of the manuscript. All authors wrote the final draft of the manuscript.

## 10. Methods

### Evo 1.5 Pretraining

We extended the pretraining of the Evo 1 model that was trained at a sequence context of 8,192 tokens with an initial learning rate of 0.003 after a warmup of 1,200 iterations; a cosine decay schedule with a maximum decay iterations of 120,000 and a minimum learning rate of 0.0003; a global batch size of 4,194,304 tokens; and 75,000 iterations (processing a total of 315 billion tokens). Other details on hyperparameters related to the model architecture and optimizer can be found in Nguyen et al. (Nguyen et al., 2024). We resumed the model’s pretraining, including all model states, optimizer states, data loading schedule, and learning rate schedule, from 75,000 iterations to 112,000 iterations (processing a total of 470 billion tokens). We refer to the model trained up to 112,000 iterations as Evo 1.5.

### Autoregressive sampling

To sample from Evo, we used a standard low-temperature autoregressive sampling algorithm (Chang and Bergen, 2023). We used the sampling code at https://github.com/evo-design/evo/ based on the reference implementation at https://github.com/togethercomputer/stripedhyena that leverages kvcaching of Transformer layers and the recurrent formulation of hyena layers (Massaroli et al., 2023; Poli et al., 2023) to achieve efficient, low-memory autoregressive generation. Parameter optimization was performed across various temperatures (0.1-1.5, increments of 0.1), top-*k* values (1-4), and top-*p* values (0.1-1.0, increments of 0.1) using the Evo 1.5 model. For each parameter combination, one hundred 1,000-nt sequences were generated from a test set of 5 gene prompts encoding 50% of a highly conserved protein. Following identification of ORFs in generated sequences using Prodigal v2.6.3 with default parameters in metagenome mode (–p meta) (Hyatt et al., 2010), generated proteins were aligned against the full-length prompt protein sequence using MAFFT (v7.526) (Katoh et al., 2002) for sequence identity calculations. For evaluating sequence degeneracy, DustMasker (version 2.14.1+galaxy2) (Morgulis et al., 2006; Camacho et al., 2009; Cock et al., 2015) was run across the full-length generations using default parameters and the proportion of masked nucleotides was calculated. Final parameters (temperature = 0.7, top-*k* = 4, top-*p*= 1.0) were selected based on maximizing sequence completion accuracy while maintaining DustMasker proportions below 0.2, a value chosen to be just slightly higher than the typical frequency of non-coding DNA in prokaryotic genomes (Rogozin et al., 2002; Troyanskaya et al., 2002).

### Sequence completion prompt compilation and evaluation

Sequences of highly-conserved genes from across prokaryotic biology were downloaded in FASTA format from NCBI GenBank (Clark et al., 2015). Selected genes included *rpoS* from *E. coli* K-12 (GenBank: NC_000913.3, coordinates 2867551-2868753), *gyrA* from *S. typhimurium* LT2 (GenBank: NC_003197.2, coordinates 2337442-2339871), and *ftsZ1* from *H. volcanii* DS2 (GenBank: NC_013967.1, coordinates 1675932-1677149). Prompts were prepared by extracting 30%, 50%, and 80% sequence lengths from the 5’ end.

Sequence completion performance was evaluated across varying prompt lengths (30%, 50%, and 80% of input sequence) using optimal sampling parameters for the Evo 1.5, Evo 1 8k (previous version of Evo trained with context length of 8,192 tokens), and Evo 1 131k (previous version of Evo trained with context length of 131,072 tokens) models (temperature = 0.7, top-*k* = 4, top-*p* = 1) (Nguyen et al., 2024). For each prompt, one hundred sequences of length 2,500 nt were generated. Prompts were subsequently appended to the start of each generated sequence and ORFs identified using Prodigal v2.6.3 with default parameters in metagenome mode (–p meta). Generated proteins were then aligned against their corresponding wild-type sequences using MAFFT (v7.526) using default parameters for sequence identity calculations.

### Operon completion prompt compilation and evaluation

Sequences encoding the *trp* operon and *modABC* operon from *E. coli* K-12 (GenBank: NC_000913.3) were downloaded in FASTA format from NCBI GenBank. For *modABC*, prompts were prepared from the full coding sequences for *modA* (coordinates 795089-795862), *modB* (coordinates 795862-796551), *modC* (coordinates 796554-797612) and *acrZ* (coordinates 794773-794922) (Clark et al., 2015). For *trp*, prompts were prepared from the full coding sequences for *trpE* (coordinates c1322946-1321384), *trpD* (coordinates c1321384-1319789), *trpC* (coordinates c1319788-1318427), *trpB* (coordinates c1318415-1317222), and *trpA* (coordinates c1317222-1316416). For bidirectional generation testing, the reverse complement of each prompt sequence was generated using Biopython (Cock et al., 2009).

Sequence completion performance was evaluated across the compiled operon completion prompts using previously identified optimal sampling parameters for Evo 1.5. For each prompt, 1000 sequences of length 2500 nt were generated. Following identification of ORFs using Prodigal, directional completion was assessed by searching *trpE*-prompted generations for *trpD*-like ORFs, *trpD* reverse complement-prompted generations for *trpE*-like ORFs, and similar pairing combinations across both *modABC* and *trp* operons. Protein sequences were then aligned against their corresponding wild-type proteins for sequence identity calculations using MAFFT (v7.526). Structural similarity was evaluated by generating protein structure predictions using Alpha Fold 3 (Abramson et al., 2024) for both generated and wild-type sequences, with structural alignments and TM-score calculations performed using TM-align (Zhang and Skolnick, 2005). Native and predicted protein structures were subsequently visualized using ChimeraX (Pettersen et al., 2021).

### Positional entropy evaluation

Per-position amino acid and nucleotide entropies were calculated from multiple sequence alignments of 500 generated and native *modB* sequences. Native *modB* sequences were fetched by querying ‘modB’ in NCBI protein, filtering by bacteria, and downloading the corresponding amino acid and nucleotide sequences in format ‘FASTA’ and ‘FASTA CDS’ respectively. Generated *modB* sequences were taken by selecting a random sample of 500 modB ORFs from the *modB* sequences generated by prompting with *modA* during the operon completion evaluation. First, nucleotide and amino acid sequences were aligned with MAFFT (v7.526) and trimmed to remove gaps appearing in >80% of sequences. For each position *i*, the Shannon entropy was then calculated as 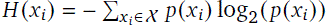 and normalized by dividing the calculated entropy by the maximum Shannon entropy (2 for nucleotides meaning all 4 bases are equally present, 4.32 for amino acids meaning all 20 standard amino acids are equally present), where *p*(*x*_*i*_) represents the frequency of each amino acid or nucleotide *x*_*i*_ at that position and *X* indicates the vocabulary.

### Toxin-antitoxin prompt compilation

Genomic loci and sequences encoding Type II toxin-antitoxin system sequences were obtained by downloading the nucleotide sequence information for all experimentally validated type II-TAs from the TADB 3.0 database (Guan et al., 2024). Using NCBI’s Entrez Programming Utilities API (EFetch from the ‘nuccore’ database using the genomic loci from TADB 3.0) (Sayers et al., 2022), the 500 bp of upstream and downstream flanking sequence were extracted for each T2TA locus. In total, for each T2TA system, six types of prompts were prepared: (1) individual toxin sequences, (2) individual antitoxin sequences, (3) the upstream context of the toxin loci, (4) the downstream context of the toxin loci, (5) the upstream context of the antitoxin loci, and (6) the downstream context of the antitoxin loci. Following successful identification of an Evo generated toxin (see section “**Evaluation of toxin activity**” below), conjugate antitoxins were subsequently generated via prompting with the generated DNA sequence encoding the toxin.

### Toxin-antitoxin sampling and filtering

To generate T2TA candidates, we first generated 57,936 sequences of 2,000 nucleotides each using Evo 1.5 (temperature = 0.7, top-*k* = 4, top-*p* = 1.0) with our compiled T2TA prompts. A multi-stage filtering pipeline was then applied to identify promising candidates. First, Prodigal was used to identify ORFs, excluding sequences containing proteins over 300 amino acids or those with only a single ORF. Next, SegMasker (version 2.14.1+galaxy2) with default parameters (Camacho et al., 2009; Cock et al., 2015) was employed to remove sequences containing low complexity regions with limited amino acid diversity.

We then assessed the protein-protein interaction potential of co-generated ORFs by co-folding all ORF pairs within each remaining generation using ESMFold. Generations were retained if they contained paired proteins with pDockQ (Basu and Wallner, 2016; Bryant et al., 2022) scores above 0.23 and high sequence dissimilarity between components. To identify novel candidates, the remaining sequences were searched against the non-redundant protein sequence database using BLAST (E-value cutoff 5 × 10^−2^), selecting for generations containing at least one component with no significant BLAST hits to known toxins or antitoxins. Final candidates were then selected based on high confidence interaction prediction using AlphaFold 2 (Jumper et al., 2021).

Following the identification of functional toxin candidates via experimental testing, the Evo-generated sequence encoding the strongest Evo-generated toxin, EvoRelE1, was used as a prompt to generate further diversified antitoxin candidates. After generating a total of 3,000 sequences from the EvoRelE1 prompt, generated ORFs were cleaned and filtered as above before being co-folded with EvoRelE1. As with the first round of generations, candidates were filtered for high pDockQ scores and low-sequence homology to known antitoxins before being evaluated for strong predicted co-folds using AlphaFold2. Final antitoxin candidates were further characterized using Foldseek web server (van Kempen et al., 2024) searches of the AlphaFold3 predicted structures (probability threshold of 0.6), blastp searches against the non-redundant protein database (E value threshold of 1), and HHpred searches (probability threshold of >90%) (Söding et al., 2005).

### Cloning of toxin and antitoxin sequences

Sequences encoding Evo-generated toxins and antitoxins were codon optimized for *E. coli* expression using IDT’s codon optimization tool and synthesized as eBlocks. The toxin plasmid backbone was prepared by PCR amplification (New England Biosciences) and gel purification (Qiagen) of pAraSpCas9+spMu (Forsberg et al., 2019) to exclude the Cas9, crRNA, tracrRNA, and gRNA sequences (**Data S1**), creating an empty arabinose-inducible vector with spectinomycin resistance. Codon-optimized toxin sequences were subsequently inserted into the modified backbone using Gibson Assembly (New England Biosciences) according to the manufacturer’s recommendations.

For antitoxin cloning, the pZE21_tetR-AcrIIA4_kanR vector (Forsberg et al., 2019) (**Data S1**) was digested with EcoRI and HindIII (New England Biosciences) and gel purified (Qiagen) to remove the AcrIIA4 sequence. Codon-optimized antitoxin sequences were then inserted into the digested vector using Gibson Assembly (New England Biosciences).

Assembled plasmids were transformed into chemically competent Stellar *E. coli* HST08 cells (Takara Biosciences), and positive clones were selected on LB agar plates containing either 50 g/mL spectinomycin (for the toxin constructs) or 50 g/mL kanamycin (for the antitoxin constructs). Plasmid sequences were then confirmed by Sanger sequencing (Elim Biosciences).

### Evaluation of toxin activity

Toxin activity was assessed through growth inhibition assays in *E. coli* NEB Turbo cells. Cells transformed with toxin constructs were first grown overnight in LB medium containing 50 g/mL spectinomycin. Cultures were then diluted 1:10 into 1 mL fresh LB supplemented with 50 g/mL spectinomycin and 2 mg/mL arabinose to induce toxin expression. After 2 hours of induction, cultures were further diluted 1:40 into 200 L of the same medium in triplicate wells of a 96-well plate. Growth was monitored using a Tecan Spark plate reader at 37°C with orbital shaking, measuring optical density (OD_600_) at 30-minute intervals over an 8-hour period. Growth curves were analyzed to evaluate the extent of toxin-mediated growth inhibition.

### Evaluation of antitoxin activity

For antitoxin evaluation, *E. coli* NEB Turbo cells were co-transformed with both toxin and antitoxin constructs and selected on LB plates containing 50 g/mL spectinomycin and 50 g/mL kanamycin. Co-transformed cells were grown overnight in LB medium containing both antibiotics. Cultures were then diluted 1:10 into 1 mL fresh LB supplemented with both antibiotics and 2 mg/mL arabinose to induce toxin expression. After 2 hours of induction, cultures were further diluted 1:40 into 200 L of the same medium in triplicate wells of a 96-well plate. Growth was monitored using a Tecan Spark plate reader at 37°C with orbital shaking, measuring optical density (OD_600_) at 30-minute intervals over an 8-hour period. Growth curves were analyzed to evaluate antitoxin-mediated rescue of growth.

Statistical comparisons were done using a one-sided Student’s *t*-test against the EvoRelE1+ eGFP negative control, with *p*-values for EvoAT1-4 being 1.61 × 10^−7^, 1.42 × 10^−7^, 2.30 × 10^−5^, and 1.69 × 10^−5^ respectively.

### Anti-CRISPR prompt compilation

Genomic sequences containing known Type II Cas9-targeting anti-CRISPR (*acr*) genes and their associated operons were obtained from previously characterized anti-CRISPR systems annotated in AcrDB. Using the Entrez API, we extracted Acr coding sequences along with 500 bp of flanking sequence both upstream and downstream of each Acr locus. For each Acr system, we prepared six types of prompts: (1) individual *acr* sequences, (2) anti-CRISPR associated (*aca*) gene sequences, (3) the upstream context of the Acr loci, (4) the downstream context of the Acr loci, (5) the upstream context of the *aca* loci, and (6) the downstream context of the *aca* loci.

### Anti-CRISPR sampling and filtering

To generate Acr candidates, we sampled a total of 3,160 sequences of 1,000 nucleotides each using Evo 1.5 with our compiled Acr prompts. A multi-stage filtering pipeline was then applied to identify promising candidates. First, Prodigal was used to identify ORFs, excluding sequences containing proteins over 200 amino acids. Next, SegMasker was employed to remove sequences containing over 50% low complexity regions or limited amino acid diversity. Sequences were subsequently folded using ESMFold to remove any candidates with very low confidence folds (pLDDT < 0.25) or no secondary structure. Candidate sequences were then evaluated using PaCRISPR, a machine learning model trained to identify potential anti-CRISPR proteins based on sequence features (Wang et al., 2020). Following guidelines established by PaCRISPR, candidates scoring above 0.5 were considered potential anti-CRISPRs, with sequences scoring above 0.75 advancing to the next step. Generated sequences classified as potential Acrs by PaCRISPR were then searched against the non-redundant protein sequence database using blastp (E-value cutoff of 1) to identify candidates with varying degrees of sequence novelty. For comparison, we also applied this filtration pipeline to evaluate sequences generated by prompting with randomly generated DNA sequences and non-Acr related genomic sequences. We then assessed the sequence diversity among the predicted Acrs by performing pairwise alignments using MAFFT (v7.526) on a set of 56 randomly selected sequences that scored above 0.75 in PaCRISPR. The resulting pairwise sequence identities were visualized using a matplotlib heatmap.

### Cloning of anti-CRISPR sequences

Sequences encoding Evo-generated anti-CRISPR proteins were codon optimized for *E. coli* expression using IDT’s codon optimization tool and synthesized as eBlocks. To generate the cloning backbone, the pZE21_tetR-AcrIIA4-Coli_kanR vector (Forsberg et al., 2019) (**Data S1**) was digested with EcoRI and HindIII (New England Biosciences) to remove the AcrIIA4 sequence and gel purified (Qiagen). Codon-optimized Acr sequences were then inserted into the digested vector using Gibson Assembly (New England Biosciences) according to the manufacturer’s recommendations. Assembled plasmids were then transformed into chemically competent Stellar *E. coli* HST08 cells (Takara Biosciences), and positive clones were selected on LB agar plates containing 50 g/mL kanamycin. Plasmid sequences were confirmed by Sanger sequencing (Elim Biosciences).

### Liquid culture assay for measuring anti-CRISPR activity

Anti-CRISPR activity was assessed through protection assays against SpCas9-mediated DNA cleavage in *E. coli*. NEB Turbo cells were first co-transformed with both the Acr expression plasmid and the CRISPR targeting plasmid containing SpCas9 and a KanR targeting guide RNA (Forsberg et al., 2019) (pAraCas9 + Sp2 + Sp6 + I-SceI, **Data S1**) and grown overnight in LB medium containing both 50 g/mL kanamycin and spectinomycin.

For liquid culture survival assays, 30 L of overnight culture was diluted into 1000 L fresh LB medium containing spectinomycin and 0.2 mg/mL arabinose to induce SpCas9 expression and deplete the kanamycin resistance plasmid. After 7 hours of growth, cultures were normalized to equal optical density and diluted 1:40 into 200 L fresh LB medium containing 50 g/mL kanamycin and spectinomycin in a 96-well plate to select for cells with active anti-CRISPR proteins. Growth was monitored using a Tecan Spark plate reader at 37°C with orbital shaking, measuring optical density (OD_600_) at 30-minute intervals over a 7-hour period. For each Acr, an uninduced control without arabinose was used to normalize growth values.

Statistical comparisons were done using a one-sided Student’s *t*-test against the random sequence negative control and AcrIIA2, with *p*-values for EvoAcr1-5 being 8.31 × 10^−6^, 6.19 × 10^−6^, 8.19 × 10^−6^, 1.43 × 10^−6^, and 8.18 × 10^−7^ against the negative control respectively and the *p*-value for EvoAcr5 against AcrIIA2 being 5.07 × 10^−4^.

### Phage plaque assay for measuring anti-CRISPR activity

For phage protection assays, *E. coli* NEB Turbo cells were co-transformed with both the Acr expression plasmid and a modified SpCas9 plasmid containing a guide RNA targeting T4(GT7) phage (SpCas9-mrh2, **Data S1**) at a 5:1 Acr:Cas9 ratio. Co-transformed cells were grown overnight in LB medium containing 50 g/mL kanamycin and spectinomycin. Cultures were then diluted 1:10 into fresh LB medium supplemented with spectinomycin, kanamycin, and 0.2 mg/mL arabinose to induce Cas9 expression. When cultures reached an OD_600_ of 0.4, 300 L was mixed into 10 mL of 0.7% soft agar containing 50 mg/mL kanamycin, 50 mg/mL spectinomycin, and 0.02 mg/mL arabinose. Plates were allowed to harden and T4(GT7) phage at a titer of 1.7 × 10^8^ PFU/mL was serially diluted 1:5 and spotted onto the bacterial lawn. Plates were then incubated overnight at 37°C before being imaged to visualize plaque formation.

### Measurement of Evo-generated protein sequence diversity

Sequence and structural diversity of generated toxin-antitoxin pairs and anti-CRISPRs were assessed through a combination of sequence and structure-based searches. Protein sequences were searched against the NCBI non-redundant protein database using blastp (E-value threshold of 1.0) to identify potential homologs. An additional search against OpenGenome was performed using MMseqs2 (version 15.6f452) with maximum sensitivity (–s 7) and other parameters set to the default values (Steinegger and Söding, 2017) to evaluate similarity to sequences in the training data. Structural similarity was evaluated by searching AlphaFold 3-predicted structures against the BFVD, AFDB-Proteome, AFDB-Swissprot, AFDB-50, BFMD, CATH50, GMGCL_ID, MG-NIFY_ESM30, and PDB100 protein databases using the Foldseek web server (https://search.foldseek. com/search) (van Kempen et al., 2024). For each generated protein, closest sequence and structural matches were identified from all three searches (BLAST, MMseqs2, and Foldseek) and evaluated to determine the degree of novelty compared to known proteins. Pairwise sequence identities were calculated from BLAST alignments, while structural similarities were quantified using Foldseek TM-scores.

### SynGenome prompt compilation

Protein sequences from prokaryotic and phage organisms were retrieved from UniProt, with all Swissprot proteins with associated coding regions and a random sample of TrEMBL proteins with associated coding regions being used as starting points for prompts (Consortium, 2024). Genomic contexts were retrieved via NCBI’s Entrez Programming Utilities (E-utilities) API, specifically using EFetch with the ‘nuccore’ database using the CDS annotations associated with the Protein Sequence accession in UniProt. For each protein’s associated genomic identifier, we constructed three API calls: one to extract the coding sequence using sequence feature coordinates, and two to extract the flanking regions by calculating positions 500 nucleotides upstream and downstream of the CDS boundaries. For CDS regions that were longer than 500 nucleotides, the prompt was derived from the final 500 nucleotides in the coding sequence. For CDS regions that were shorter than 500 nucleotides, the CDS sequence was trimmed to the nearest 100 bp. Additionally, we generated reverse complement sequences for each region using the Biopython Seq module, resulting in six distinct prompts per protein: upstream, coding sequence (CDS), downstream, and their respective reverse complements. Associated functional annotations including Gene Ontology (GO) terms (Ashburner et al., 2000; The Gene Ontology Consortium et al., 2023), species names, and InterPro domains (Blum et al., 2024) were retrieved from UniProt’s REST API for each protein using their UniProt accession numbers and linked to their corresponding prompts.

### SynGenome sampling

Sequences were generated using Evo 1.5 with optimized sampling parameters (temperature = 0.7, top-*k* = 4, top-*p* = 1.0). For each prompt type, we generated 2 sequences of 5,000 nucleotides in length, yielding a total database size of over 120 billion base pairs. Generation was performed in parallel across multiple compute nodes to facilitate large-scale sequence production.

### Data filtering of SynGenome sequences

Generated sequences underwent a multi-step filtering pipeline to remove low-complexity regions while preserving biologically plausible features. Initial validation removed any invalid characters from the nucleotide sequences. These sequences were then processed using DustMasker (NCBI BLAST+ v2.16.0) with a masking level of 30 (–level 30) and FASTA output format (–outfmt fasta) to identify low-complexity regions. Following DustMasker processing, we implemented two additional filtering steps: removal of successive 100 nucleotide chunks from the sequence end if they contained more than 40% masked bases, and elimination of any continuous masked regions longer than 800 nucleotides. Sequences were excluded from the final database based on several criteria: length below 100 nucleotides, masked base content exceeding 80% in sequences shorter than 2,000 nucleotides, complete masking of all bases, or empty/NaN values. The sequences passing these filtering criteria were converted to uppercase and retained in the final database.

### Prediction of SynGenome ORFs

Open reading frames were identified in the filtered sequences using Prodigal v2.6.3 with default parameters and metagenomic prediction mode (–p meta). Predictions were initially refined by excluding sequences shorter than 40 amino acids or longer than 1200 amino acids and sequences with incomplete protein sequences. Following basic filtration, low-complexity regions were identified using NCBI SegMasker (window size 15, locut 1.8, hicut 3.4), with sequences containing >20% masked regions being excluded. Additional complexity filters removed sequences with fewer than 12 unique amino acids or highly repetitive k-mer patterns (k=3-10, threshold >40% coverage). This multi-step filtering process ensured the retention of high-quality protein predictions while removing potentially spurious or low-complexity sequences.

### Creation of SynGenome website

The SynGenome website was implemented with HTML, CSS, and JavaScript to provide a searchable web interface at https://evodesign.org/syngenome/. The interface allows users to query sequences using protein names, UniProt IDs, InterPro domains, species names, or GO terms. The database was structured to maintain associations between generated sequences and their corresponding prompt annotations. The raw SynGenome data is hosted on Hugging Face for public access.

### Creation of SynGenome UMAP and Leiden Clusters

To generate the UMAP, first, a random sample of 50,000 sequences encoding at least one ORF with prompts derived from the CDS sequence was extracted from SynGenome. This random sample was subsequently filtered to remove generations with < 40% or > 60% GC content, resulting in a final set of 36,762 generations. Embeddings were generated for both the prompt and corresponding generated sequences by extracting activations from the MLP layer 3 in the 6th hyena block of Evo 1.5’s architecture before being mean pooled along the sequence dimension to create fixed-length representations. These high-dimensional embeddings were subsequently reduced to two dimensions using UMAP (n_neighbors=15, min_dist=0.1). Sequences were colored according to their percent identity to sequences in the training data, calculated using MMseqs2 against a database of prokaryotic genomes used in model training. These high-dimensional embeddings were reduced to two dimensions using UMAP with ScanPy (version 1.10.3) (Wolf et al., 2018) default parameters. Sequences were colored according to their percent identity to sequences in the training data, calculated using MMseqs2 against OpenGenome. Graph-based clustering was performed using the default Leiden algorithm implemented in Scanpy (version 1.10.3). The resolution parameter was optimized by evaluating cluster stability across resolutions from 0.1 to 0.5, measuring both the number of clusters and coefficient of variation of cluster sizes. A final resolution of 0.2 was selected for clustering based on these metrics.

### Comparison of SynGenome to native prokaryotic sequences

To evaluate how representative SynGenome was of natural prokaryotic sequences, first, a random sample of 36,762 5,000-nt generations was taken from OpenGenome to match the number of sequences used in the representative sample of SynGenome. Following the procedure used for prediction of ORFs in SynGenome (see section “Prediction of SynGenome ORFs”), Prodigal was used to identify potential ORFs in the sampled sequences and minimal filtering was applied to remove clearly incorrectly called sequences. Length distributions of predicted SynGenome ORFs were compared to those from the random OpenGenome sample. Codon usage bias of the prompts and SynGenome generations were analyzed by calculating the total frequency of each codon across all prompt ORFs and all generated ORFs and normalizing the frequencies to the total number of codons per dataset. Protein family domains were identified in both datasets using HMMER v3.3.2 hmmscan (hmmer.org) against the Pfam-A database (Paysan-Lafosse et al., 2024) with default parameters (E-value cutoff 0.01). Domain frequencies were compared between datasets using Fisher’s exact test with Benjamini-Hochberg multiple testing correction (minimum occurrence threshold of 10 domains in both datasets). The relationship between domain frequencies was visualized using a log-scale scatter plot, with points colored by absolute log_2_ fold change between datasets. Correlation between domain frequencies was assessed using the Pearson correlation coefficient calculated using the pearsonr function in SciPy (Virtanen et al., 2020). The distribution of domain occurrence frequencies was analyzed by binning domains found in the scanned SynGenome and OpenGenome proteins into log-scale bins (2^*n*^ occurrences) and visualizing the count of unique domains per frequency bin for each.

## Supplementary Material

**Figure S1.**
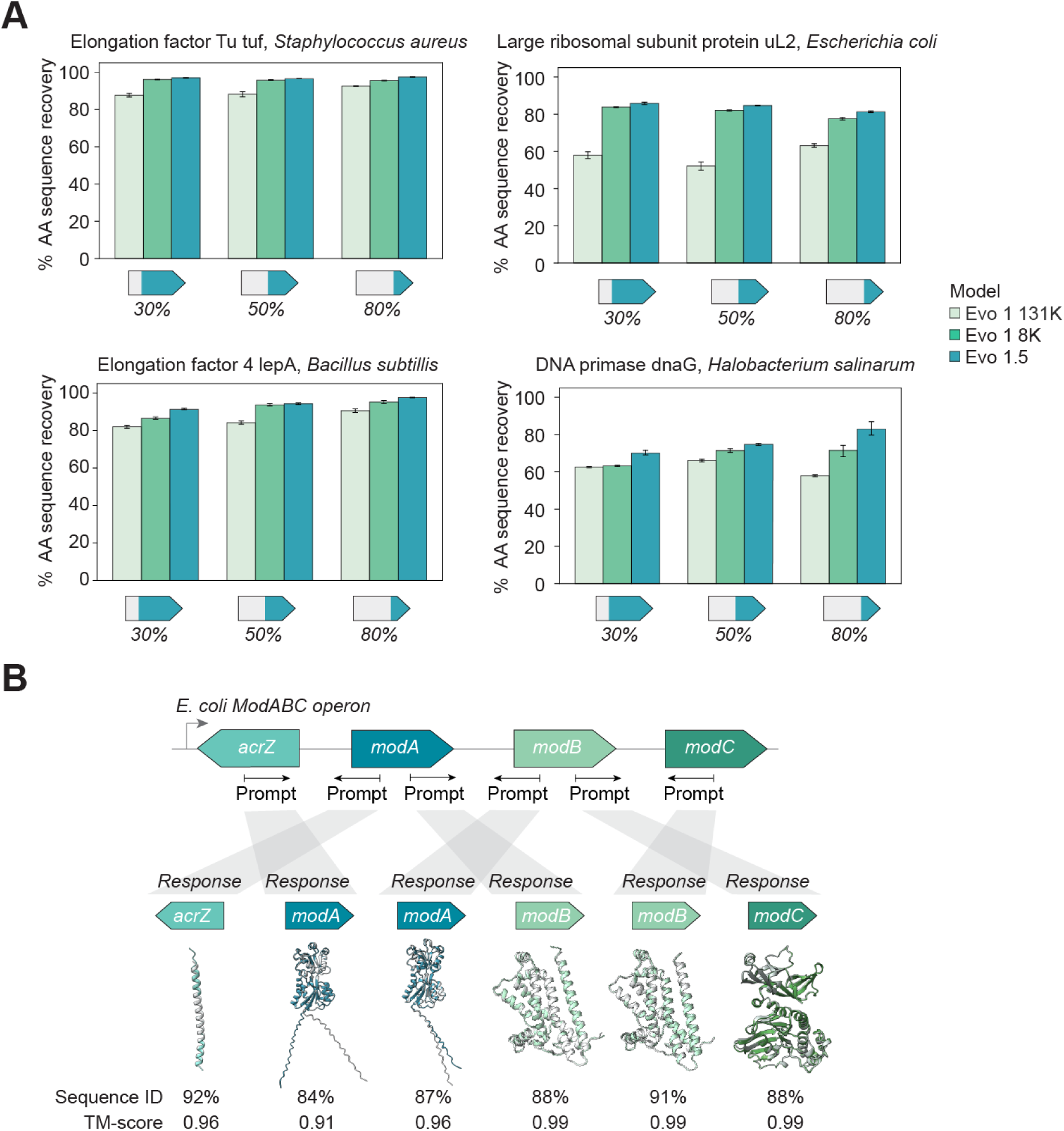
| Additional gene and operon completion evaluations. **(A)** Bar graph showing gene completion of conserved genes *EF-Tu*, *uL2*, *lepA*, and *dnaG* highlights the ability of Evo to understand genomic localization from prompts alone. AA, amino acid; Bar height, mean; error bars, standard error; *n* = 100. **(B)** Bidirectional completion of conserved E. coli modABC operon gene sequences demonstrates high sequence identity and structural conservation across the operon.

**Figure S2.**
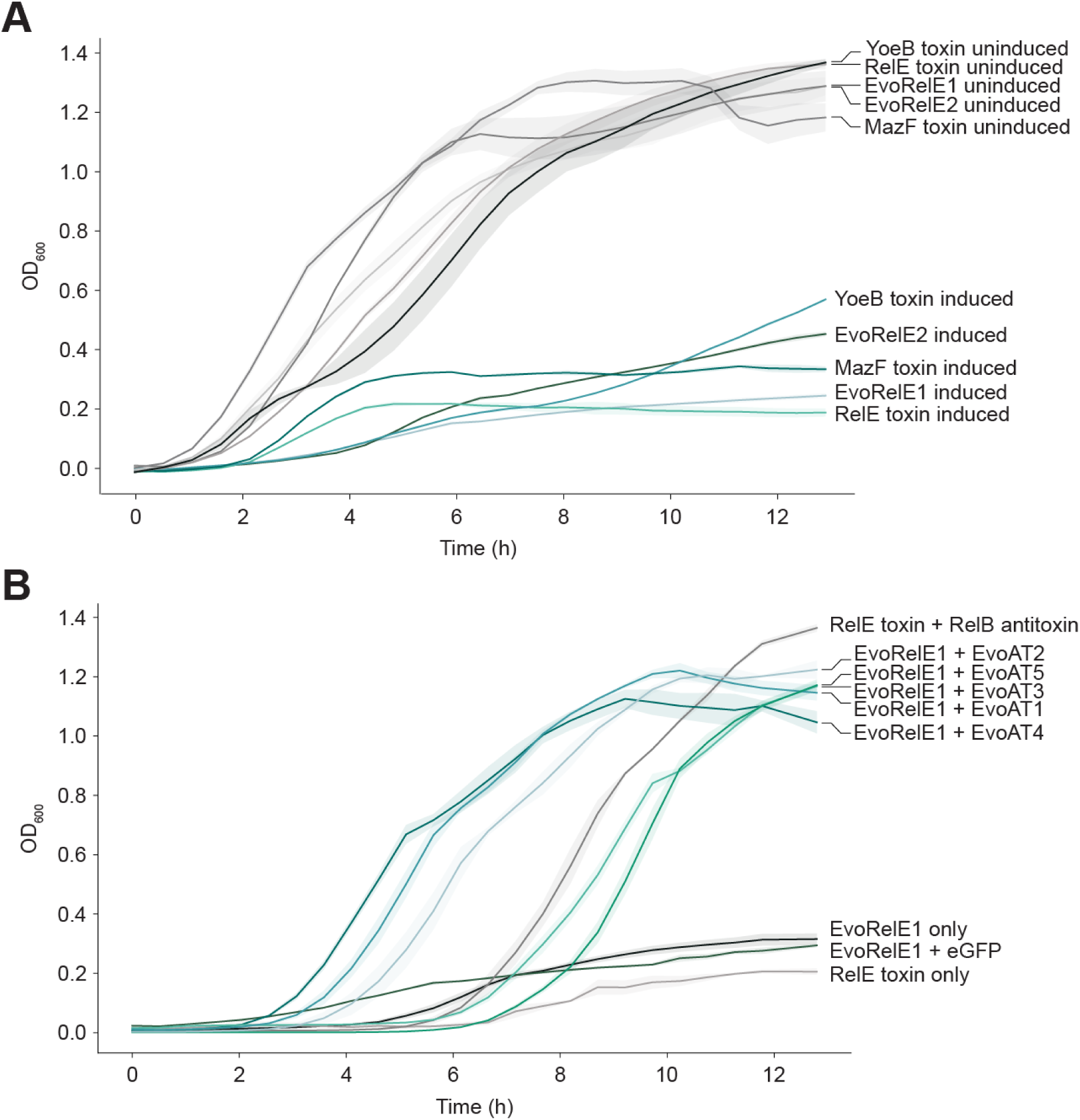
| Growth curves for toxin and antitoxin growth inhibition and rescue assays. **(A)** Growth curves show growth arrest in E. coli when toxins EvoRelE1, EvoRelE2, and all native toxins are induced. **(B)** Growth curves show growth rescue when EvoAT1-5 are present in combination with EvoRelE1 and growth arrest when only EvoRelE1 is present.

**Figure S3.**
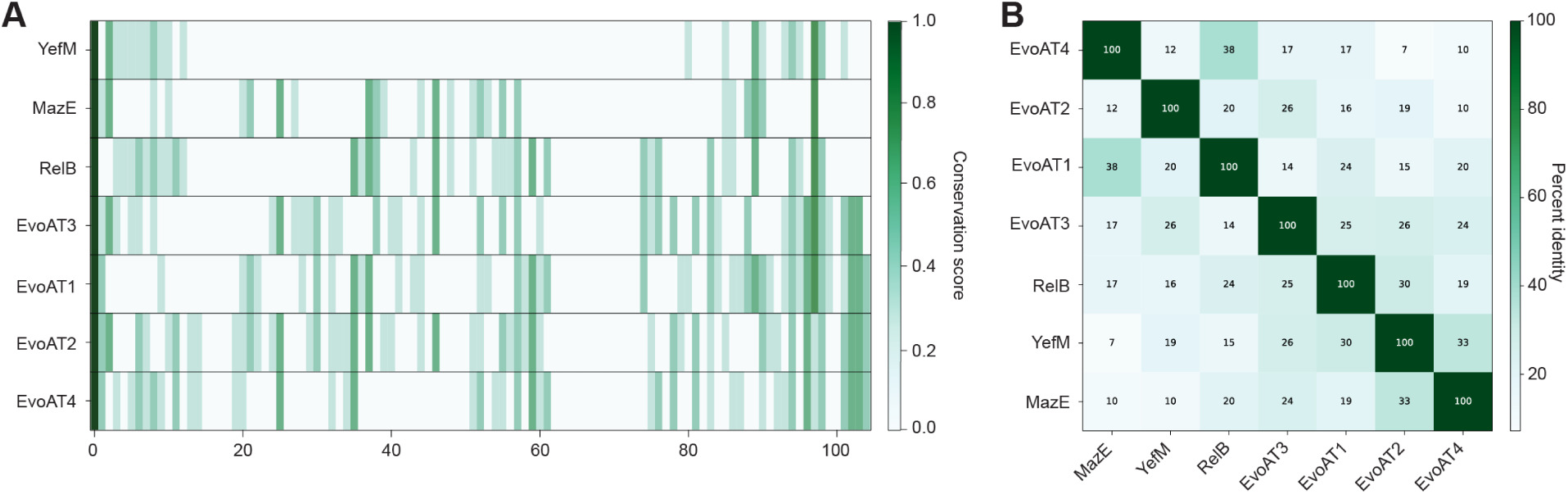
| Alignment of Evo antitoxins with native antitoxins. **(A)** Homology analysis of aligned amino acid sequences for EvoAT1-4 against *E. coli* MazE, YefM, and RelB, with darker green indicating higher conservation. **(B)** Percent identity matrix for EvoAT1-4 against against E. coli MazE, YefM, and RelB, with darker green representing higher percent identity. EvoAT1-4 share limited homology with wild-type antitoxins.

**Figure S4.**
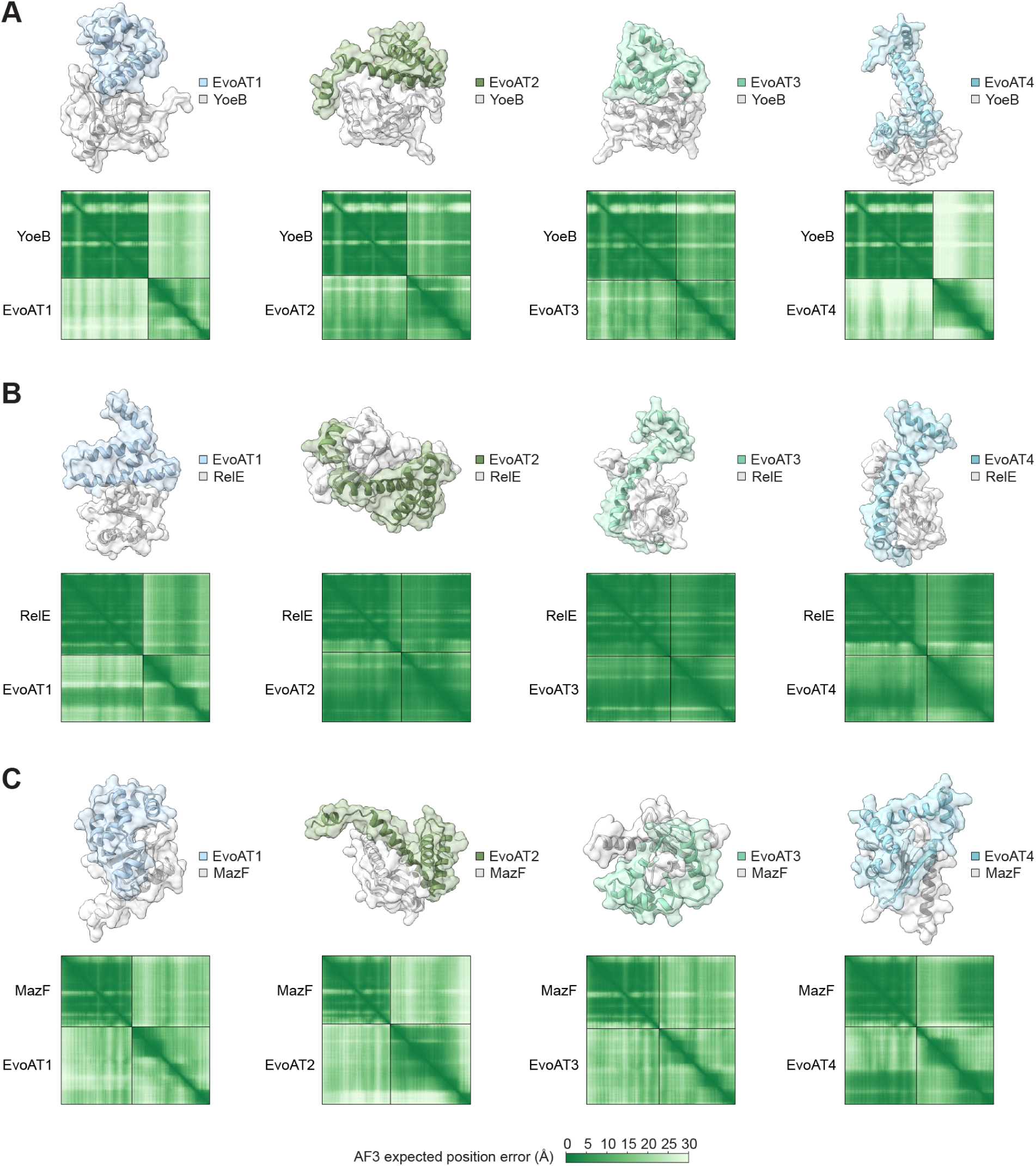
| AlphaFold 3 structure predictions for Evo antitoxins in complex with native toxins. **(A)** Evo antitoxins in complex with YoeB **(A)**, RelE **(B)**, and MazF **(C)**.

**Figure S5.**
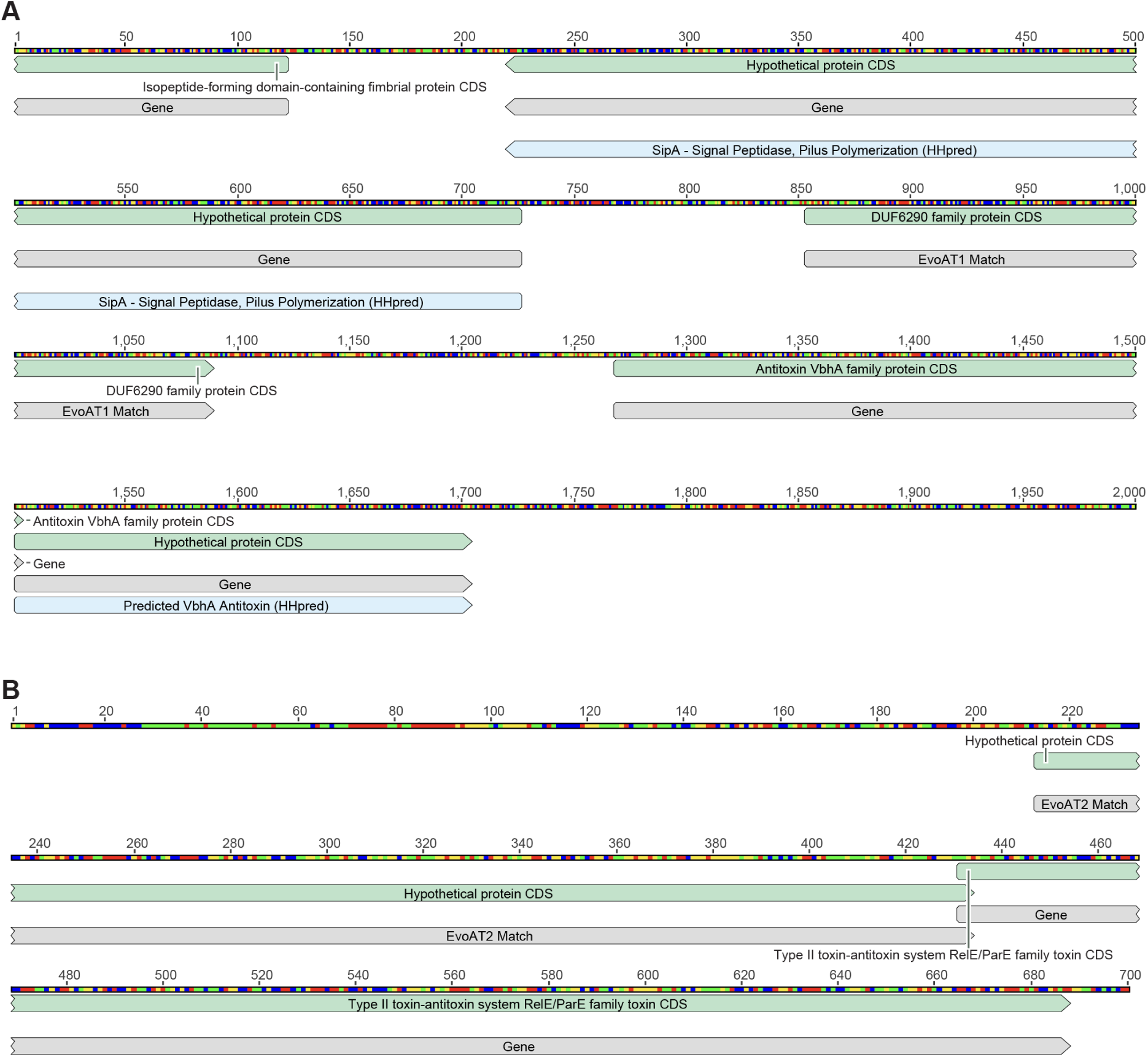
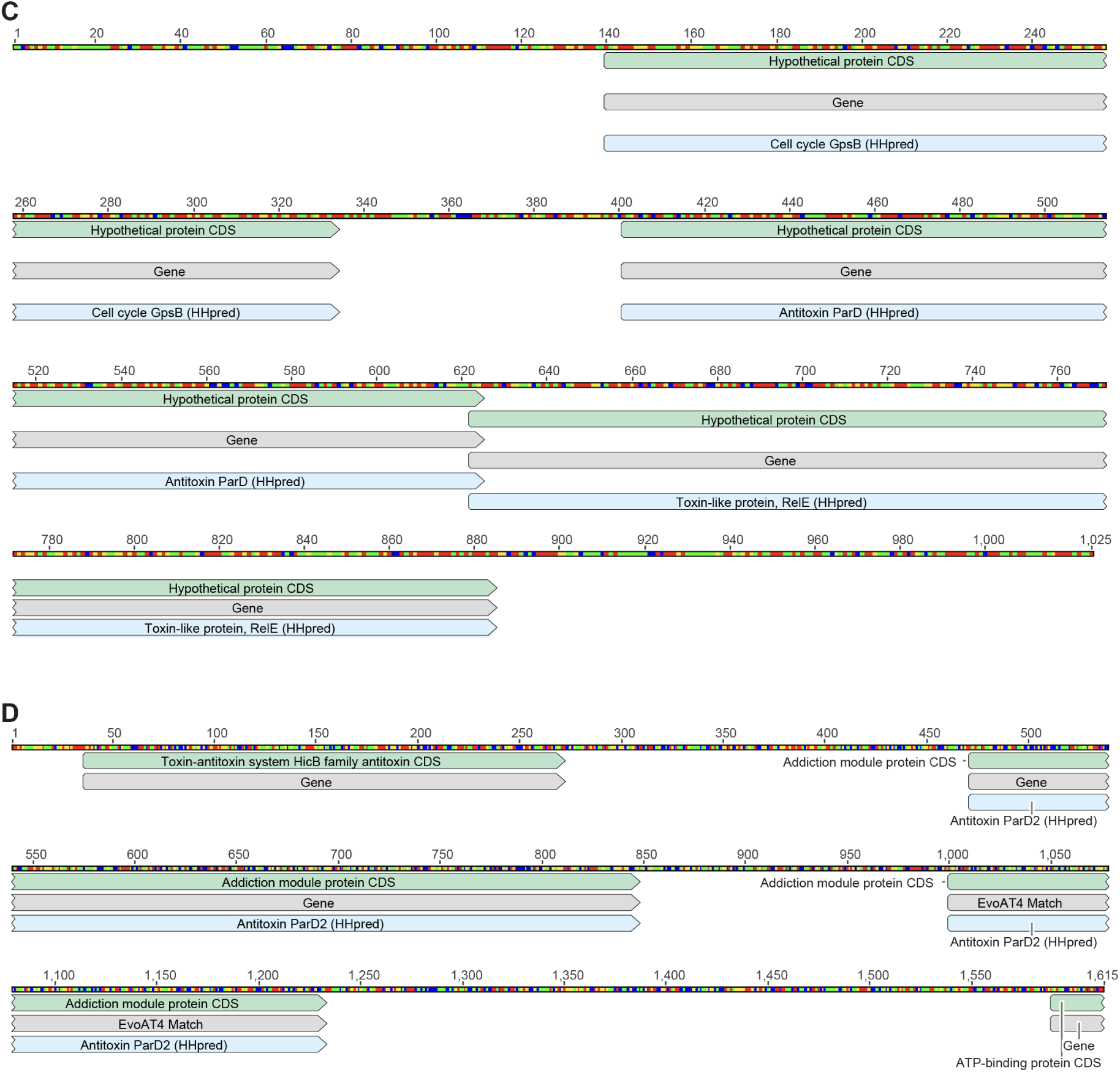
| Operonic context of Evo antitoxin sequence matches. **(A-D)** Genes known and predicted to be related to toxin-antitoxin systems can be found in the immediate vicinity of the closest sequence identity matches for EvoAT1-4.

**Figure S6.**
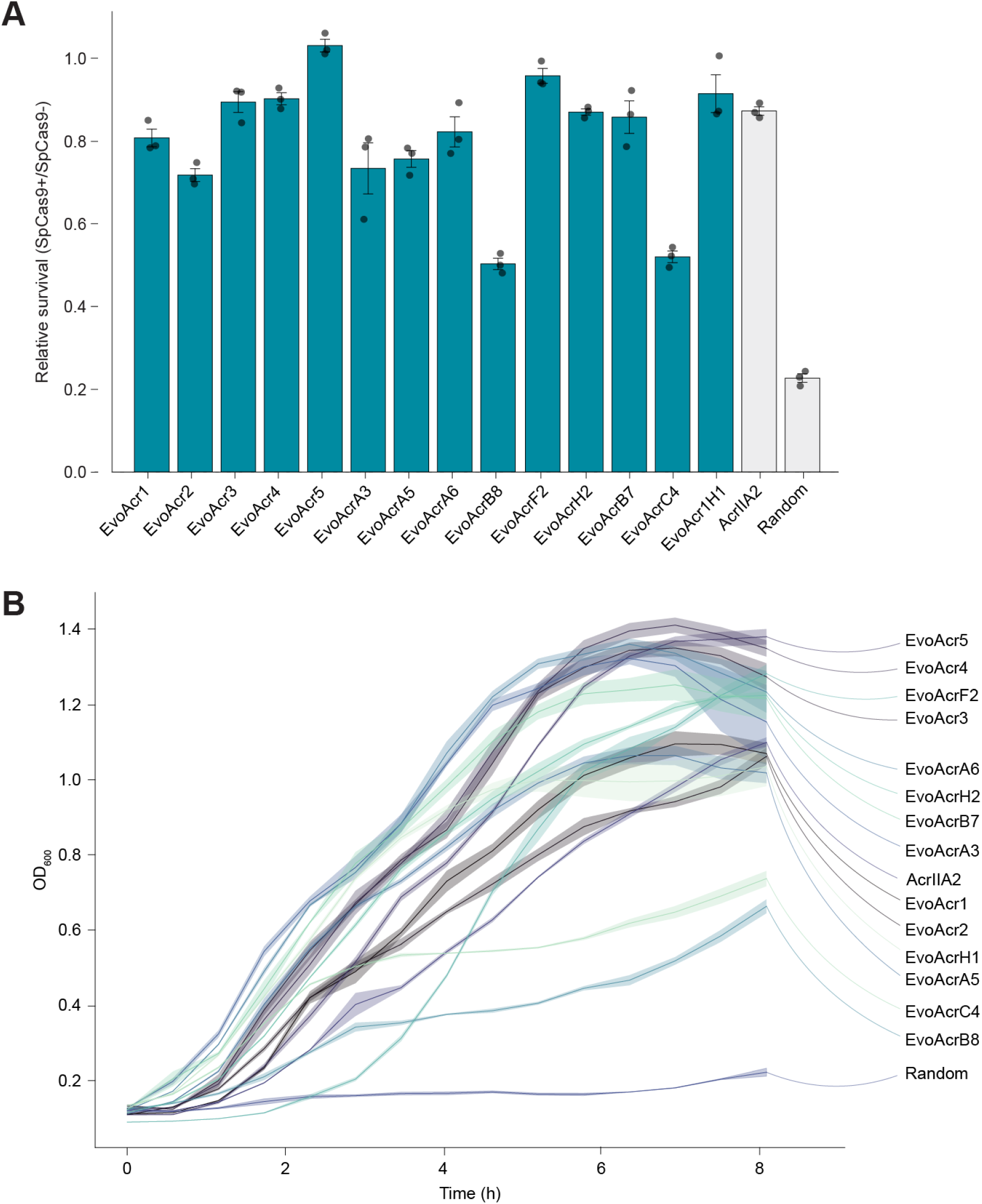
| Growth curves for Acr protection assay. **(A)** Bar plot showing relative survival rates for all successful Evo-generated Acrs. Bar height, mean; error bars, standard error; circles, biological replicate values; *n* = 3. **(B)** Growth curves show growth rescue when functional EvoAcrs are present with KanR-targeted SpCas9 and cell death when only non-functional sequences are present.

**Figure S7.**
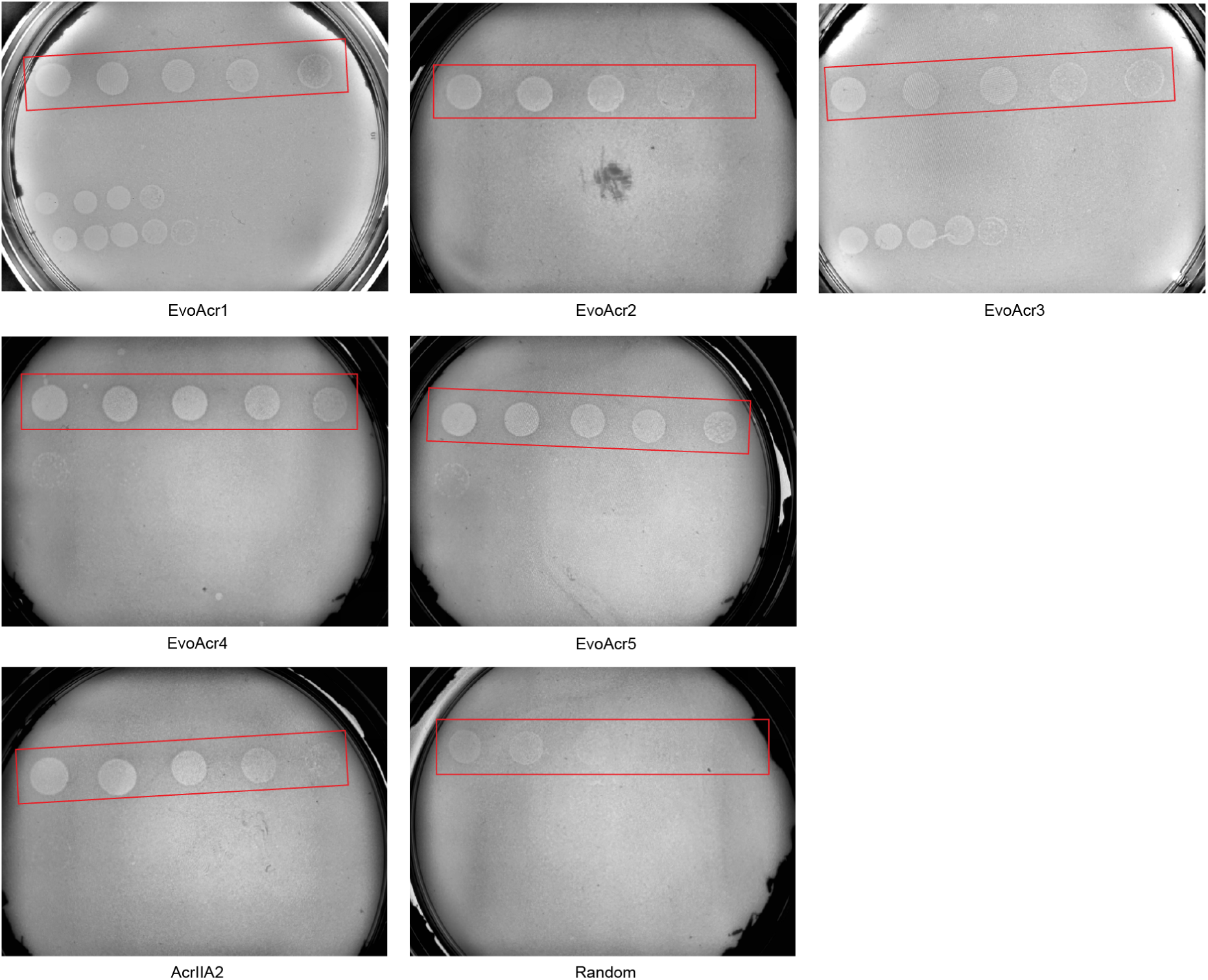
| Phage plaque images for Acr protection assay. Uncropped phage plaque images depicting for-mation of plaques in response to EvoAcr1-5, AcrIIA2, and a random sequence.

**Figure S8.**
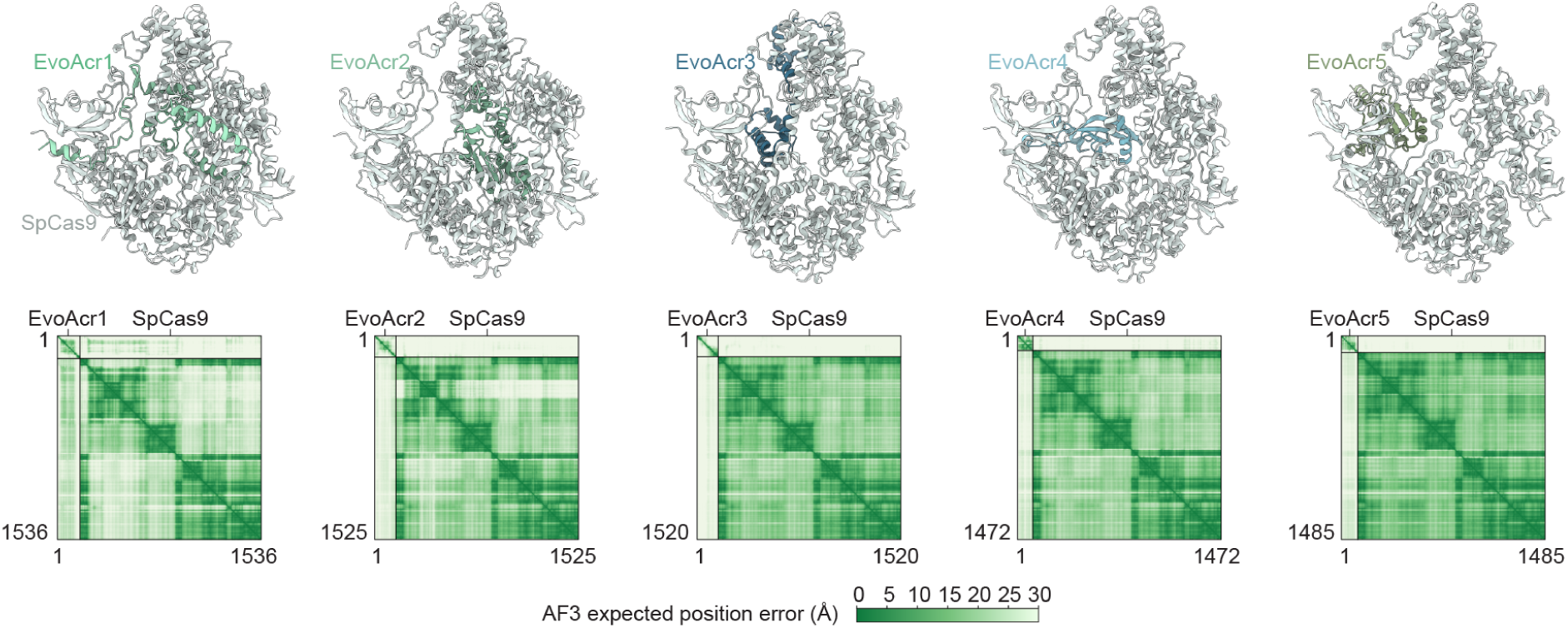
| AlphaFold 3 structure predictions for Evo Acrs in complex with SpCas9. AlphaFold 3 structure predictions of Evo Acrs in complex with SpCas9 (top) and their corresponding predicted aligned error (PAE) plots (bottom).

**Figure S9.**
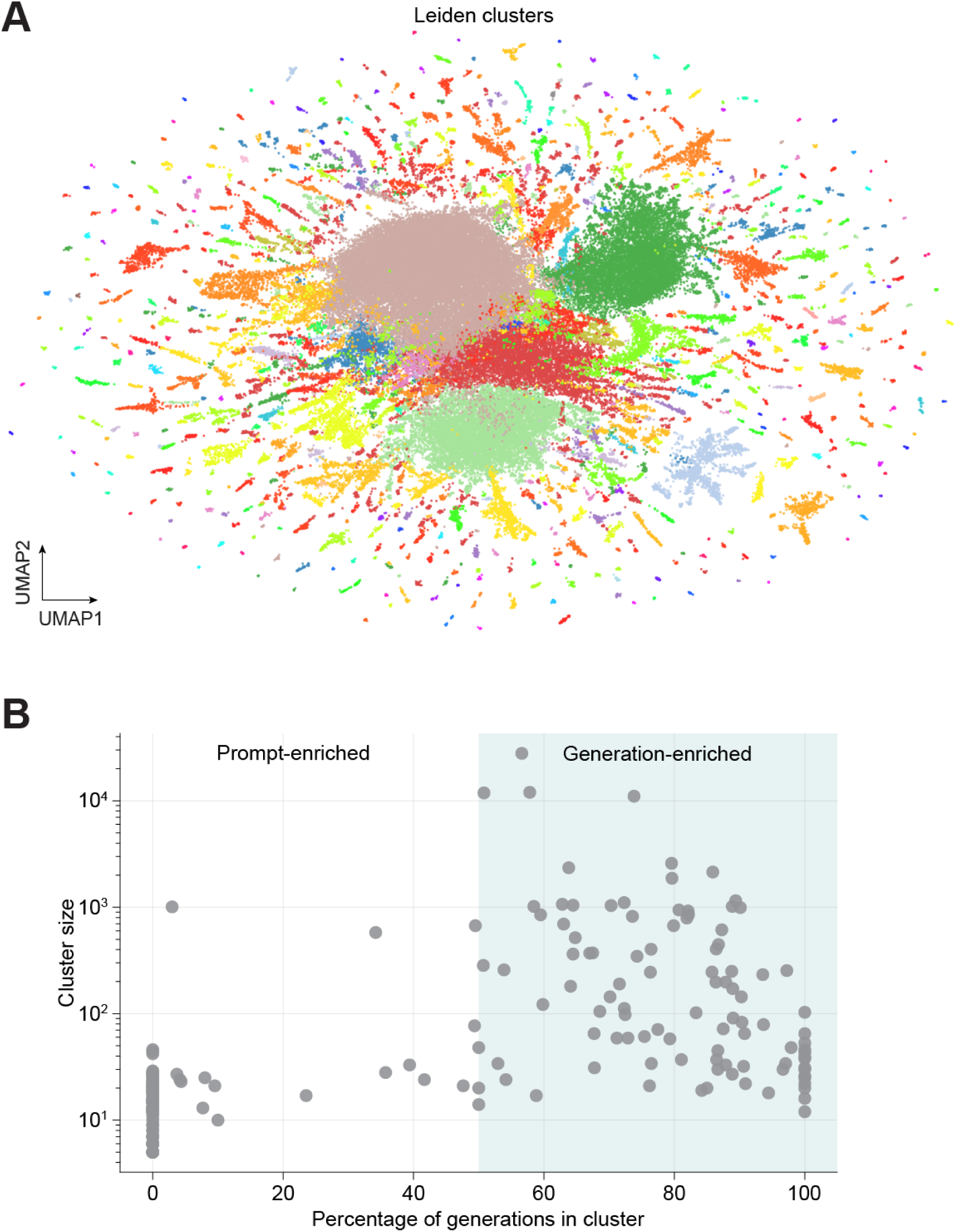
| Extended Leiden cluster analysis of SynGenome. **(A)** UMAP of Leiden clusters of Evo embeddings of prompt sequences and generated sequences. **(B)** Distribution of prompts and generations across Leiden clusters.

**Figure S10.**
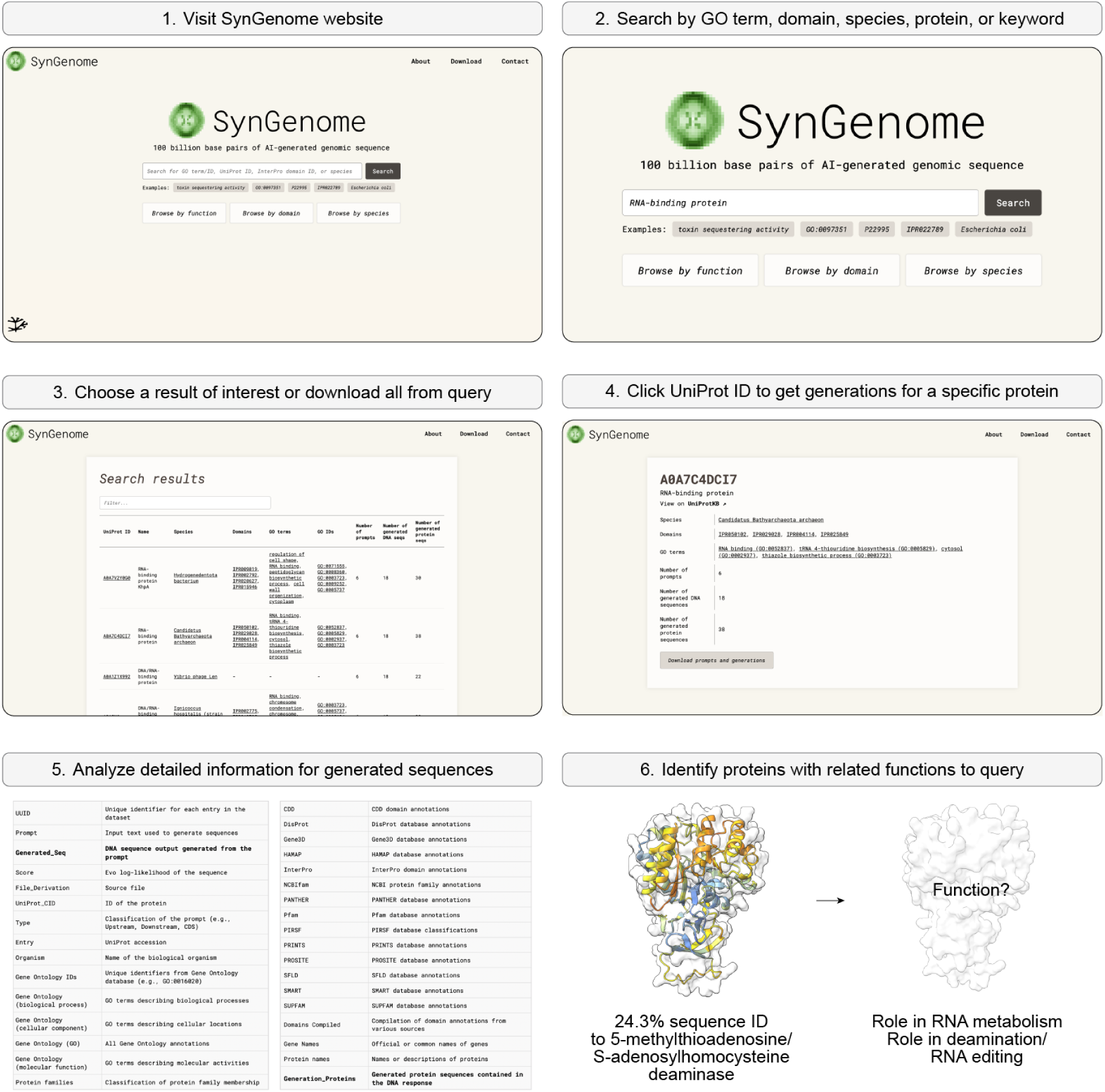
| Example walkthrough of the SynGenome web-interface workflow. SynGenome can easily be queried with a domain, function, species, or keyword of interest to find relevant generated sequences, which can then be downloaded to identify potential functionally-related protein candidates. We demonstrate the applicability of SynGenome by showing its generation of a potential RNA modification enzyme in response to a RNA binding protein-derived prompt.

